# The globular C1q receptor is required for epidermal growth factor receptor signaling during *Candida albicans* infection

**DOI:** 10.1101/2021.05.25.445718

**Authors:** Quynh T. Phan, Jianfeng Lin, Norma V. Solis, Michael Eng, Marc Swidergall, Feng Wang, Shan Li, Sarah L. Gaffen, Tsui-Fen Chou, Scott G. Filler

## Abstract

During oropharyngeal candidiasis, *Candida albicans* activates the epidermal growth factor receptor (EGFR), which induces oral epithelial cells to both endocytose the fungus and synthesize proinflammatory mediators that orchestrate the host immune response. To elucidate the signaling pathways that are stimulated when *C. albicans* interacts with EGFR, we analyzed the proteins that associate with EGFR when *C. albicans* infects human oral epithelial cells. We identified 1214 proteins that were associated with EGFR in *C. albicans*-infected cells. We investigated the function of seven of these proteins that either showed increased association with EGFR in response to *C. albicans* or that mediated the interaction of other microbial pathogens with epithelial cells. Among these proteins, EGFR was found to associate with WW domain-binding protein 2, toll-interacting protein, interferon-induced transmembrane protein 3, and the globular C1q receptor (gC1qR) in viable epithelial cells. Each of these proteins was required for maximal endocytosis of *C. albicans* and they all regulated fungal-induced production of IL-1β and/or IL-8, either positively or negatively. gC1qR functioned as a key coreceptor with EGFR. Interacting with the *C. albicans* Als3 invasin, gC1qR was required for the fungus to stimulate both EGFR and the ephrin type-A receptor 2. The combination of gC1qR and EGFR was necessary for maximal endocytosis of *C. albicans* and secretion of IL-1β, IL-8, and GM-CSF. Thus, this work provides an atlas of proteins that associate with EGFR and identifies several that play a central role in the response of human oral epithelial cells to *C. albicans* infection.

**IMPORTANCE:** Oral epithelial cells play a key role in the pathogenesis of oropharyngeal candidiasis. In addition to being target host cells for *C. albicans* adherence and invasion, they secrete proinflammatory cytokines and chemokines that recruit T cells and activated phagocytes to foci of infection. It is known that *C. albicans* activates EGFR on oral epithelial cells, which induces these cells to endocytose the organism and stimulates them to secrete proinflammatory mediators. To elucidate the EGFR signaling pathways that govern these responses, we analyzed the epithelial cell proteins that associate with EGFR in *C. albicans-infected* epithelial cells. We identified four proteins that physically associate with EGFR and that regulate different aspects of the epithelial response to *C. albicans.* One of these is gC1qR, which is required for *C. albicans* to activate EGFR, induce endocytosis, and stimulate the secretion of proinflammatory mediators, indicating that gC1qR functions as a key co-receptor with EGFR.

## INTRODUCTION

*Candida albicans* grows as a harmless commensal in the oral cavity of at least 50% of normal adults (1). However, when either local or systemic immune defenses are weakened, *C. albicans* can proliferate and cause oropharyngeal candidiasis (2). This disease causes substantial morbidity in patients with HIV/AIDS, dentures, organ transplantation, cancer, diabetes, and xerostomia (3–6). Although most patients with their first episode of oropharyngeal candidiasis readily respond to treatment with antifungal agents such as fluconazole, patients with recurrent disease are at risk for developing infection with an azole-resistant strain (7, 8).

During oropharyngeal candidiasis, *C. albicans* invades the epithelial cell lining of the oropharynx, stimulating a strong proinflammatory host response that is driven by IL-17 and IL-22 signaling that is initiated from innate lymphocytes and subsequently from adaptive immune response (9, 10). When *C. albicans* adheres to an epithelial cell, it interacts with and activates multiple epithelial cell receptors, including the ephrin type-A receptor 2 (EphA2), E-cadherin, the platelet-derived growth factor receptor BB, HER2, and the epidermal growth factor receptor (EGFR) (11–13). Among these receptors, EGFR plays a central role in triggering the epithelial cell endocytosis of *C. albicans* and stimulating a proinflammatory response. EGFR interacts with the ephrin type-A receptor 2 (EphA2), HER2, E-cadherin, src family kinases, and the aryl hydrocarbon receptor, triggering changes in the actin cytoskeleton that lead to the endocytosis of the fungus (11, 13, 14). EGFR also interacts with EphA2 to stimulate epithelial cells to secrete proinflammatory mediators, including defensins, IL-1α, IL-1β, CXCL8/IL-8, and CCL20 (11, 15, 16). The defensins have direct candidacidal activity, while the cytokines and chemokines recruit phagocytes to the focus of infection and enhance their capacity to kill the invading fungus.

Because of the key role of EGFR in mediating the oral epithelial cell response to *C. albicans,* we sought to obtain a more comprehensive view of the spectrum of host cell proteins that interact with this receptor. Using a proteomics approach, we found that EGFR associates with 1214 epithelial cell proteins in *C. albicans-infected* cells. Of these proteins, 13 had increased association with EGFR in the *C. albicans-infected* cells relative to uninfected cells. In intact epithelial cells, four proteins associated with EGFR in the vicinity of *C. albicans* hyphae: WW domain-binding protein 2 (WBP2) which governs EGFR expression and signaling in cancer cells, toll-interacting protein (TOLLlIP) which is a negative regulator of toll-like receptor signaling, interferon-induced transmembrane protein 3 (IFITM3) which is an antiviral protein, and the globular C1q receptor (gC1qR) which is a multifunctional protein that interacts with a variety serum components. We interrogated these in the context of *C. albicans* infection of human oral epithelial cells, and found that each of these proteins is required for maximal endocytosis of *C. albicans.* Moreover, they all regulate the production of innate cytokines such as IL-1β and/or IL-8, either positively or negatively. Additionally, gC1qR functions as a key coreceptor that is required for *C. albicans* to stimulate EGFR and to induce endocytosis and an epithelial cell proinflammatory response.

## RESULTS

### EGFR associates with multiple proteins that mediate endocytosis and govern actin dynamics

To identify epithelial cell proteins that associate with EGFR, we infected the OKF6/TERT-2 oral epithelial cell line (17) with *C. albicans* yeast for 90 min, cross-linked the proteins with formaldehyde and then performed immunoprecipitation of whole cell lysates with an anti-EGFR antibody. The proteins that were associated with EGFR were then identified by LC-MS/MS. As controls, we processed in parallel uninfected epithelial cells as well as epithelial cells incubated with epidermal growth factor (EGF) for 5 min. In total, 1214 proteins were associated with EGFR in all three samples of epithelial cells infected with *C. albicans* (Table S1) and 1278 were associated with EGFR in both samples of epithelial cells that had been incubated with EGF (Table S2). The majority of these proteins were constitutively associated with EGFR. However, 13 proteins showed at least a 2-fold increase in association with EGFR in all three samples of *C. albicans-*infected cells relative to uninfected cells (Table 1) and 37 had increased association with EGFR in the EGF-exposed cells (Table S3). Among the proteins that had increased association with EGFR in response to *C. albicans* infection, four associated with EGFR only in cells exposed to *C. albicans*, but not EGF. Among these were: WW domain-binding protein 2 (WBP2), guanylate binding protein family member 6 (GPB6), subunit J of eukaryotic translation initiation factor 3, and calcineurin subunit B type 1 (Table 1). Only three proteins were found to associate with EGFR in cells exposed to EGF but not *C. albicans.* One was EGF itself and the others were galectin-9B and ERBB receptor feedback inhibitor 1 (Table S3). Collectively, these results suggest that while the profiles of proteins that associate with EGFR in response to *C. albicans* and EGF are similar, some proteins are uniquely recruited in response to *C. albicans*.

**Table 1.** List of proteins that had increased association with EGFR in response to infection with *C. albicans.* Shown are individual results from 3 biological replicates for epithelial cells infected with *C. albicans* and 2 biological replicates from epithelial cells incubated with EGF. Values are the ratio of the amount of protein detected in cells exposed to *C. albicans* or EGF relative to unstimulated control cells. Inf indicates proteins that were undetectable in unstimulated control cells.

| Accession Number | Gene | Protein | *C. albicans*-1 | *C. albicans*-2 | *C. albicans*-3 | EGF-1 | EGF-2 |
| --- | --- | --- | --- | --- | --- | --- | --- |
| K7EIJ0 | WBP2 | WW domain-binding protein 2 | Inf | Inf | Inf | - | - |
| B4DRS8 | GBP6 | cDNA FLJ54753, highly similar to guanylate binding protein family, member 6 | Inf | Inf | Inf | - | - |
| B4DUI3 | EIF3J | Eukaryotic translation initiation factor 3 subunit J | 2.63 | Inf | Inf | - | - |
| F6U1T9 | PPP3R1 | Calcineurin subunit B type 1 | Inf | 2.15 | Inf | 0.00 | - |
| Q9H0E2 | TOLLIP | Toll-interacting protein | Inf | Inf | Inf | Inf | - |
| P35354 | PTGS2 | Prostaglandin G/H synthase 2 | 3.29 | Inf | Inf | - | Inf |
| Q15075 | EEA1 | Early endosome antigen 1 | 15.80 | 12.72 | 3.11 | 6.38 | 1.83 |
| P32926 | DSG3 | Desmoglein-3 | 3.30 | 5.99 | 2.08 | 0.61 | 0.76 |
| Q01628 | IFITM3 | Interferon-induced transmembrane protein 3 | 3.17 | 3.18 | 2.68 | 0.00 | 0.91 |
| Q14165 | MLEC | Malectin | 2.10 | 2.46 | 2.57 | 1.45 | 1.61 |
| P02787 | TF | Serotransferrin | 4.06 | 2.05 | 2.24 | 2.30 | 1.32 |
| P21281 | ATP6V1B2 | V-type proton ATPase subunit B, brain isoform | 4.10 | 7.36 | Inf | 4.03 | Inf |
| P36543 | ATP6V1E1 | V-type proton ATPase subunit E 1 | 2.47 | Inf | 2.26 | Inf | 1.94 |

Analysis of proteins constitutively associated with EGFR provided insight into the mechanisms by which EGFR induces epithelial cells to endocytose *C. albicans.* We found that EGFR associated with EphA2, E-cadherin, and HER2 (Table 2), consistent with our previous findings that EGFR functions in the same pathways as these three receptors (11, 13, 18). The src family kinases are known to activate EGFR (14), and we found that two members of this family, Lyn and Yes associate with this receptor (Table 2). Receptors are sometimes located in specific microdomains on the plasma membrane. It was notable that EGFR was associated with caveolin-1, caveolin-2, flotillin-1, flotillin-2, arf6, and RhoA (Table 2), which are components of lipid rafts (19, 20).

**Table 2.**
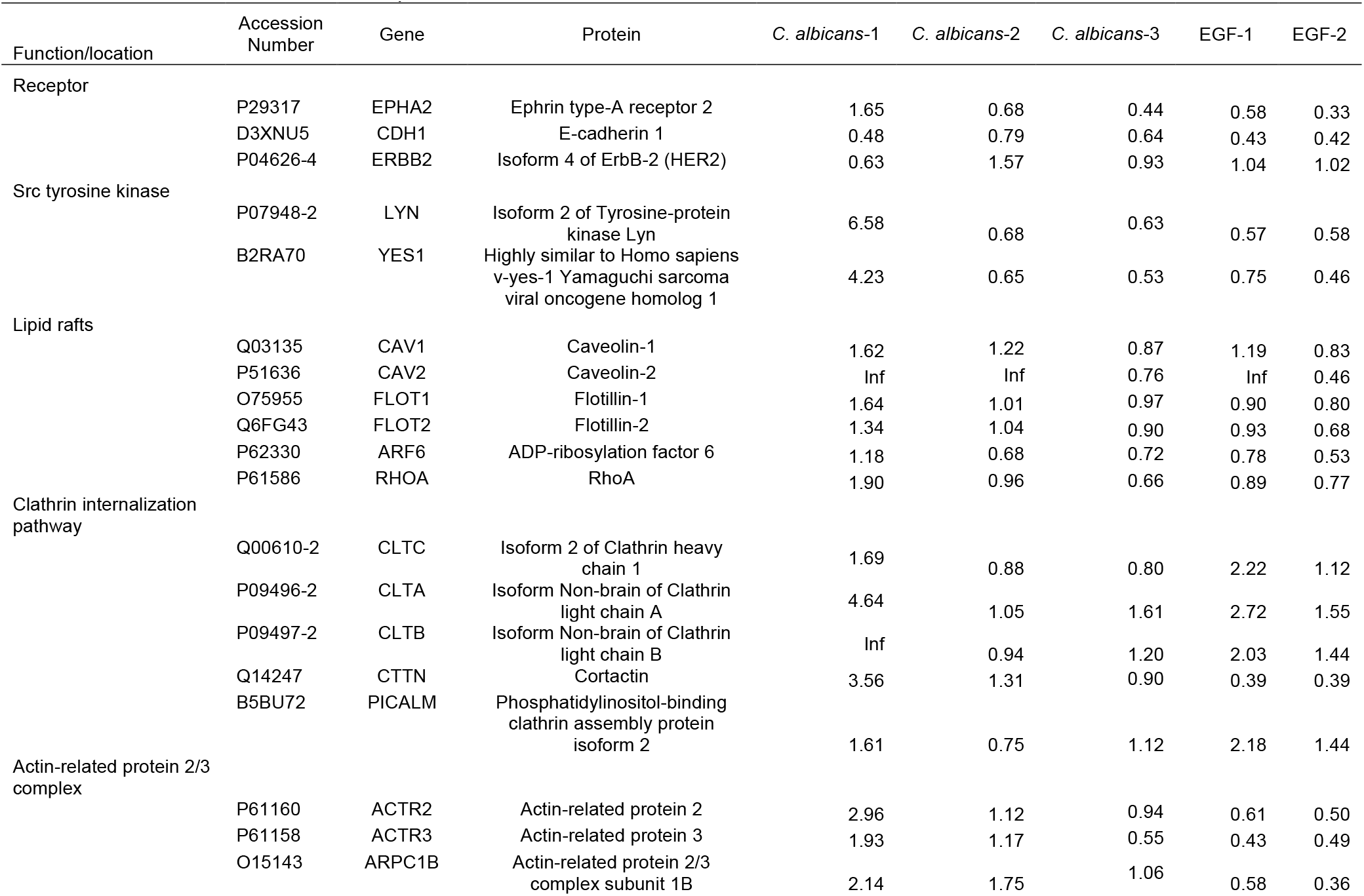
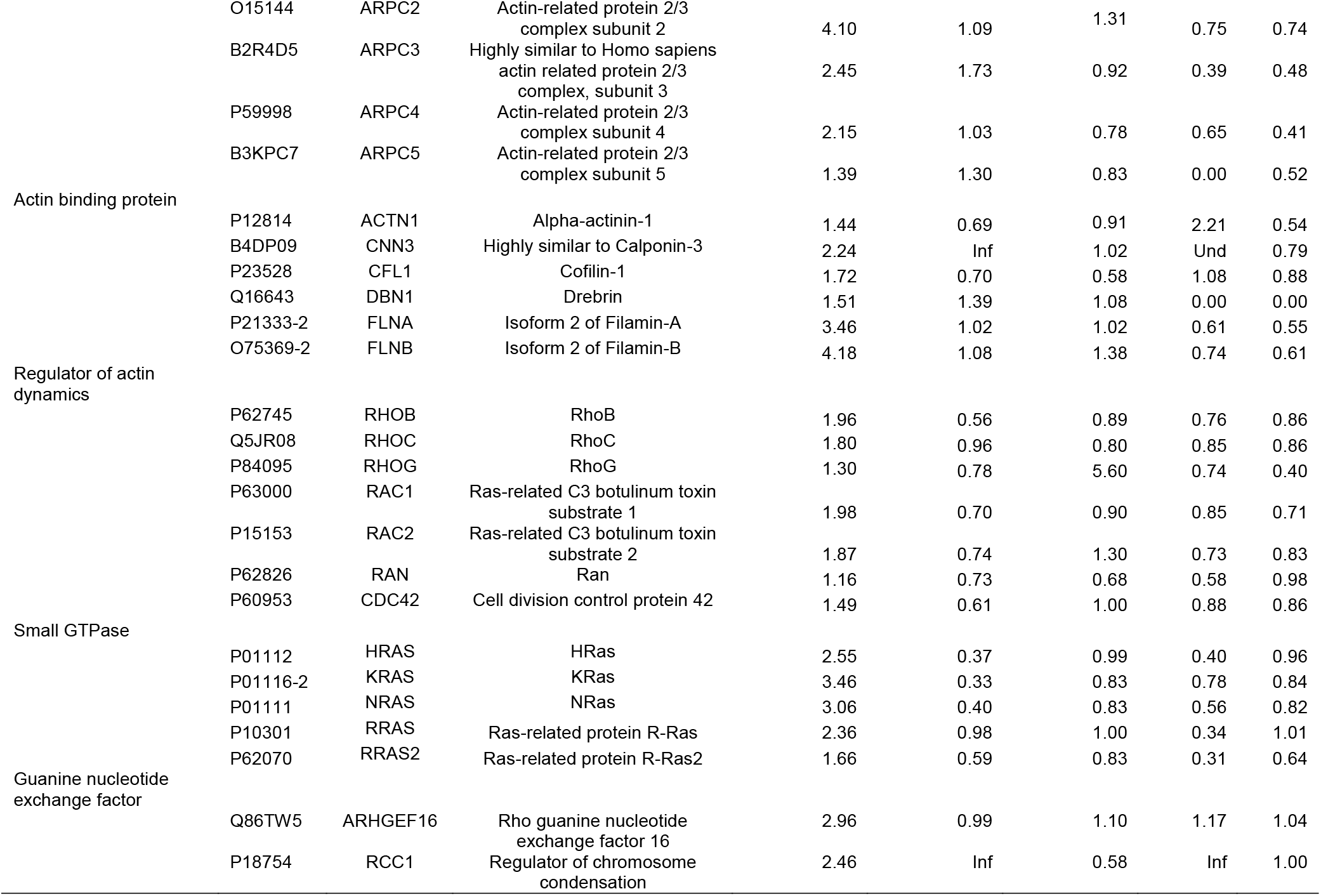
List of proteins that had constitutive association with EGFR in response to infection with *C. albicans.* Shown are individual results from 3 biological replicates for epithelial cells infected with *C. albicans* and 2 biological replicates from epithelial cells incubated with EGF. Values are the ratio of the amount of protein detected in cells exposed to *C. albicans* or EGF relative to unstimulated control cells. Inf indicates proteins that were undetectable in unstimulated control cells.

The endocytosis of *C. albicans* is also mediated in part by the clathrin and cortactin internalization pathway (21). Consistent with this mechanism, the clathrin heavy and light chains and cortactin were found to associate with EGFR (Table 2). Also associated with EGFR was the phosphatidylinositol-binding clathrin assembly protein, an adapter protein that is required for proper clathrin function (20). Activation of the clathrin pathway induces rearrangement of actin filaments, leading to the formation of pseudopods that engulf the fungus, leading to the formation of an endocytic vacuole (22). We found that EGFR associated with all seven members of the actin-related protein 2/3 (arp2/3) complex, including actin-related proteins 2 and 3, and actin-related protein complex subunits 1B, 2, 3, 4, 5 (Table 2). These proteins play a key role in organizing actin filaments during clathrin-mediated endocytosis and the formation of pseudopods (23, 24).

EGFR was also associated with numerous actin binding proteins, including α-actinin, calponin 3, cofilin-1, drebrin, and fascin (Table 2). The association of EGFR with filamin A and B, which link actin to membrane glycoproteins (25) suggests that these both may connect EGFR with actin and its associated proteins. Actin dynamics are known to be regulated by small GTPases (26), and we found that EGFR was associated with RhoA, RhoB, RhoC, RhoG, RAC1, RAC2, Ran, and CDC42 (Table 2). EGFR associated with additional small GTPases, including HRas, KRas, NRas, R-Ras, R-Ras2 and 28 different Rabs (Table 2). Most small GTPases require guanine nucleotide-exchange factors (GEFs) for activation (27), and EFGR associated with guanine nucleotide exchange factor 16 (neuroblastoma, ephexin 4) and regulator of chromosome condensation 1 (RCC1) (Table 2). Collectively, these results indicate that EGFR has extensive interactions with the actin cytoskeleton and its regulatory proteins that likely induce the endocytosis of *C. albicans* hyphae and secretion of proinflammatory mediators when EGFR is activated.

### The EGFR-associated proteins WBP2, TOLLIP, IFITM3, and gC1qR govern the epithelial cell response to *C. albicans*

Among the proteins that were predicted by the proteomics data to have increased association with EGFR in response to *C. albicans,* we selected six for additional study: WBP2, guanylate binding protein 6 (GBP6), TOLLIP, early endosome antigen 1 (EEA1), desmoglein-3 (DSG3), and IFITM3. These proteins were chosen because of their potential roles in governing the endocytosis of *C. albicans* and the secretion of proinflammatory mediators. We also investigated the globular C1q receptor (gC1qR), which was constitutively associated with EGFR. Our rationale was that gC1qR is known to function as a receptor for *L. monocytogenes,* and there are significant similarities between the mechanisms of host cell invasion by this bacterium and *C. albicans* (18, 21, 28–30).

To verify that the selected proteins were associated with EGFR in intact oral epithelial cells and to determine their subcellular location with respect to *C. albicans* cells, we used a proximity ligation assay (16). This assay forms a flourescent spot where two proteins are located within 40 nm of one another (31). All seven proteins were associated with EGFR in both infected and uninfected epithelial cells, but only WBP2, TOLLIP, IFITM3, and gC1qR accumulated with EGFR in the vicinity of *C. albicans* hyphae (Fig. 1 and Fig. S1). The accumulation of WBP2, TOLLIP, IFITM3, and gC1qR around *C. albicans* suggested that these proteins might govern the epithelial cell response to the fungus.

**Fig. 1.**
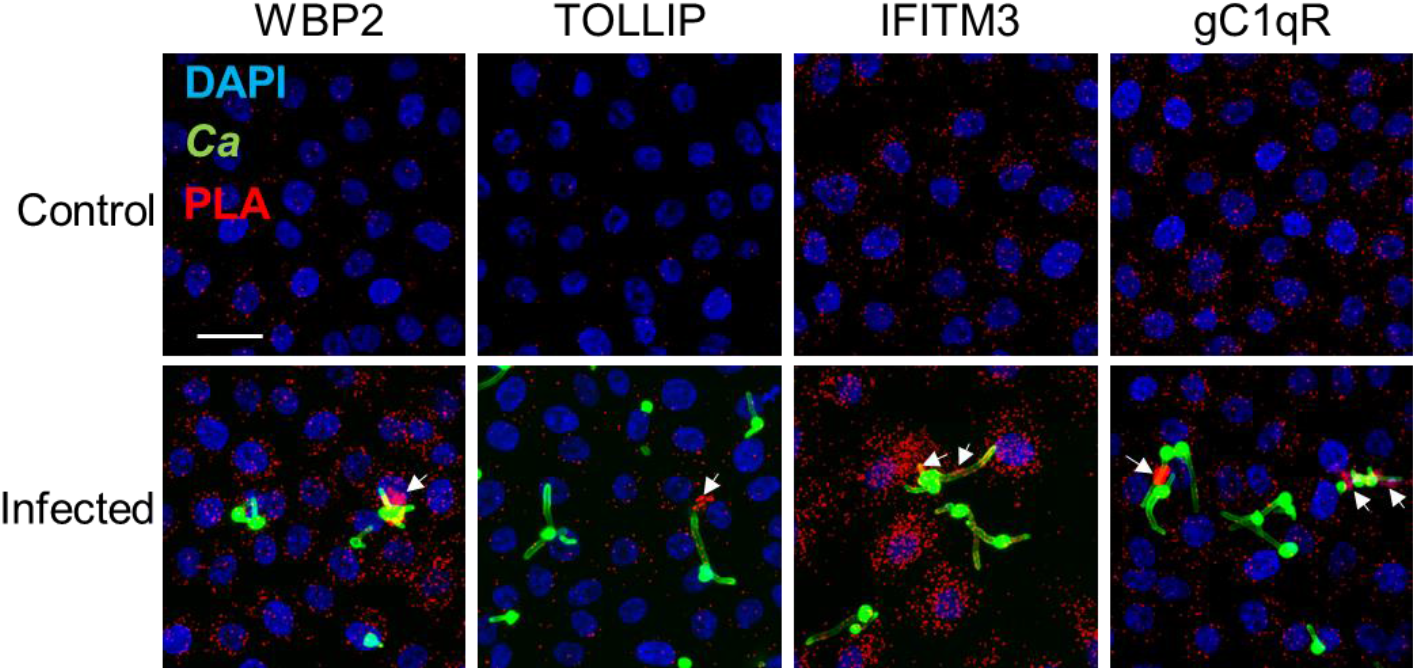
Proximity ligation assays showing the physical association of the epidermal growth factor receptor (EGFR) with WW domain-binding protein 2 (WBP2), toll-interacting protein (TOLLIP), interferon-induced transmembrane protein 3 (IFITM3), and the globular C1q receptor (gC1qR) in the OKF6/TERT-2 oral epithelial cell line. The epithelial cells were incubated with either medium alone (top) or infected with *C. albicans* (bottom) for 90 min. Red spots indicate the regions where the indicated proteins associate with EGFR. Arrows indicate the accumulation of the proteins around *C. albicans* hyphae. Results are representative of three independent experiments. Scale bar 25 μm.

To investigate this hypothesis, we used siRNA to knock down the levels of each these proteins in oral epithelial cells. We then measured *C. albicans* adherence to epithelial cells and subsequent endocytosis using a standard differential fluorescence assay (18, 22). We also analyzed the secretion of IL-1β and IL-8 by ELISA. As a control, we used siRNA to knock down EGFR in the epithelial cells. Consistent with previous results (13, 16), knockdown of EGFR modestly reduced the number of cell-associated *C. albicans* cells and decreased the number of endocytosed organisms by almost 50% (Fig. 2A). Reduction of EGFR also inhibited *C. albicans-* induced secretion of IL-1β and Il-8.

**Fig. 2.**
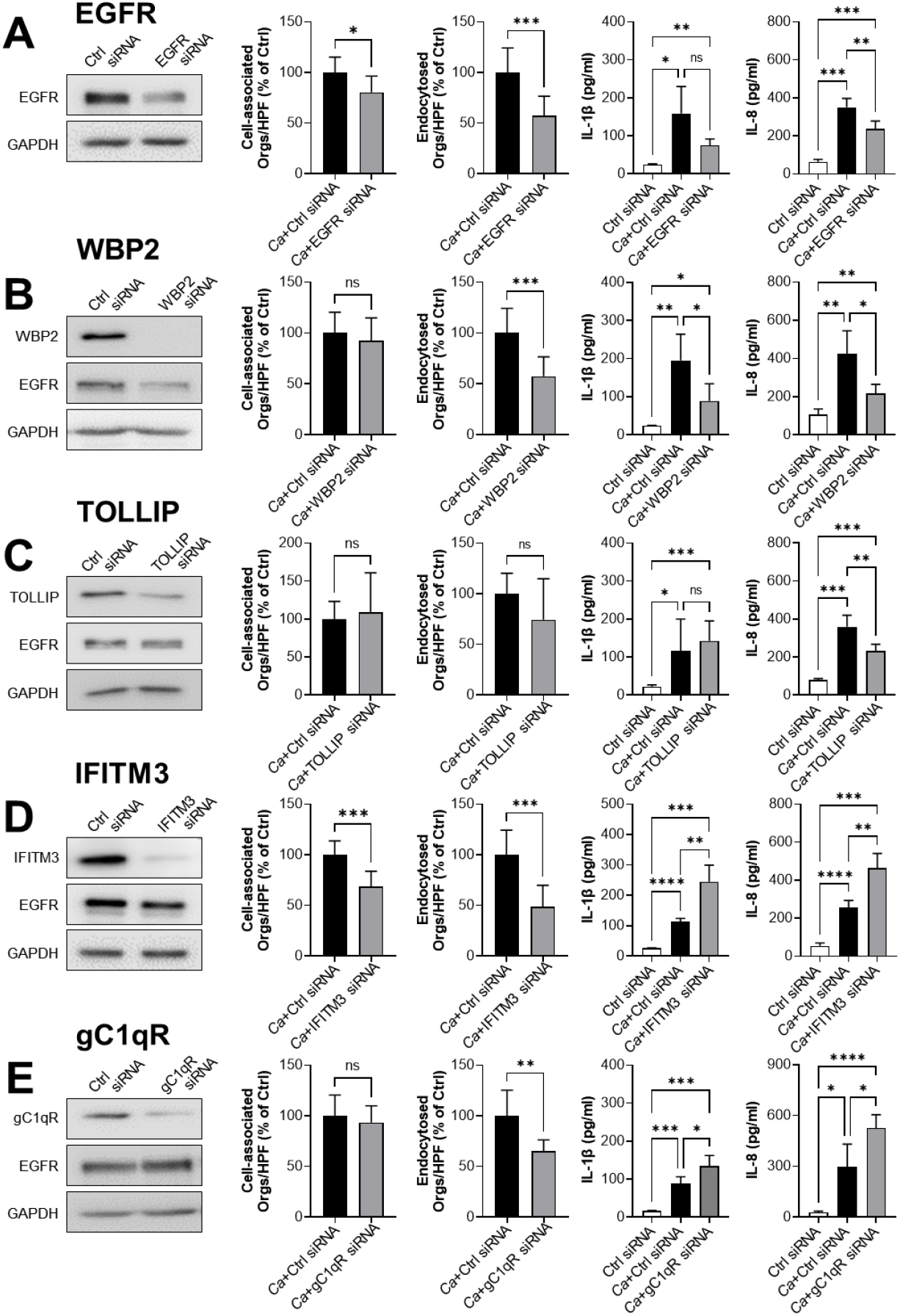
Functional analysis of proteins that interact with EGFR. Oral epithelial cells were transfected with EGFR (A), WBP2 (B), TOLLIP (C), IFITM3 (D), and gC1qR (E) siRNA. For each siRNA, the extent of protein knockdown and its effects on the number of cell-associated organisms, the number of endocytosed organisms, IL-1β secretion, and IL-8 secretion were determined. The graphs show the mean ± SD of three independent experiments, each performed in triplicate. The data were analyzed using one-way analysis of variance with Dunnett’s test for multiple comparisons. Ca, *C. albicans;* Ctrl, control; NS, not significant; \**P* < 0.05; \*\**P* < 0.01; \*\*\**P* < 0.001; \*\*\*\**P* < 0.0001.

WBP2 is a multifunctional protein that has mainly been studied in the context of breast cancer. Functioning as an adapter protein and transcriptional co-activator, WBP2 is required for maximal EGFR expression and for the normal activity of the PI3K/Akt signaling pathway (32). WBP2 also links JNK to Wnt signaling (32). Both PI3k/Akt and JNK have been shown to govern the response of oral epithelial cells to *C. albicans* (33, 34). As shown, siRNA knockdown of WBP2 reduced total EGFR levels in oral epithelial cells, leading to a decrease in *C. albicans* endocytosis and a reduction in IL-1β and IL-8 production (Fig. 2B). Thus, knockdown of WBP2 largely phenocopies the knockdown of EGFR, and is required for normal cellular EGFR levels in oral epithelial cells.

TOLLIP is a membrane-associated endocytic adapter protein that is a negative regulator of the innate immune response. TOLLIP inhibits signaling by STAT1, toll-like receptor, and IL-1β (35–37), but was not previously known to associate with EGFR. Knockdown of TOLLIP inhibited secretion of IL-8 but had no impact on the adherence or endocytosis of *C. albicans* or the secretion of IL-1β (Fig. 2C). These results suggest that TOLLIP is a positive regulator of epithelial cell IL-8 secretion in response to *C. albicans* infection.

IFITM3 plays a key role in the host defense against viral infections by binding to virus particles and shuttling them to lysosomes for degradation (38). By a similar process, IFITM3 also enhances the degradation of activated EGFR (38). Knockdown of IFITM3 in oral epithelial cells had paradoxical effects. Although loss of IFITM3 inhibited the adherence and endocytosis of *C. albicans,* it stimulated *C. albicans-induced* secretion of IL-1β and IL-8 (Fig. 2D), thereby dissociating the process of endocytosis from cytokine production.

gC1qR (HABP1/p32) is present in the mitochondrial matrix where is involved in oxidative phosphorylation (39). Although gC1qR lacks a transmembrane sequence, it is also expressed on the cell surface where it functions as a receptor for C1q, high-molecular weight kininogen, factor XII, vitronectin, and hyaluronic acid (40–43). gC1qR It also acts as a host cell receptor for multiple pathogens including *L. monocytogenes, Staphylococcus aureus,* and *Plasmodium falciparum* (28, 44, 45). In carcinoma cells, inhibition of gC1qR with a monoclonal antibody is known to reduce EGFR phosphorylation and block stimulation of migration and lamellipodia formation in response to EGF (46). Knockdown of gC1qR inhibited the endocytosis of *C. albicans* but stimulated the secretion of IL-1β and IL-8 (Fig. 2E), suggesting that gC1qR may play a key role in the activation of EGFR by *C. albicans*.

### Surface-exposed gC1qR is required for EGFR-mediated endocytosis of *C. albicans*

Based on the above results, we focused on gC1qR for in-depth study. Because gC1qR is expressed both on the cell surface and intracellularly, the gC1qR siRNA likely reduced the levels of both cell surface and intracellular gC1qR. To test whether the inhibition of just surface-exposed gC1qR altered the epithelial cell response to *C. albicans*, we evaluated two different anti-gC1qR monoclonal antibodies, 60.11 and 74.5.2 that bind to different domains of gC1qR (47). Both antibodies decreased the endocytosis of *C. albicans* (Fig. 3A), and antibody 74.5.2 slightly reduced the number of cell-associated organisms (Fig. S2A). Also, treating the epithelial cells with both monoclonal antibodies together did not result in a further inhibition of endocytosis. Thus, surface-exposed gC1qR is required for maximal endocytosis of *C. albicans* by oral epithelial cells.

**Fig. 3.**
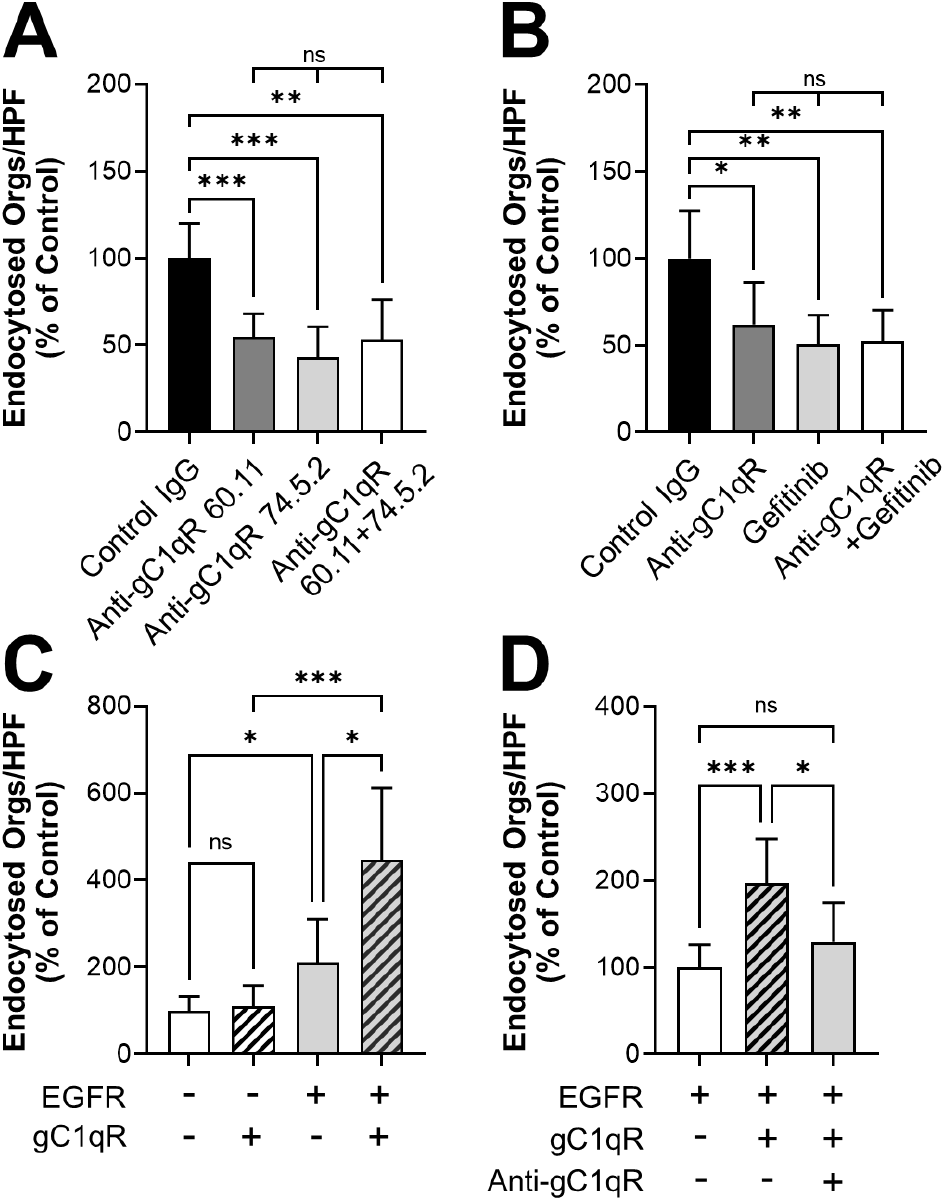
Surface-expressed gC1qR mediates the endocytosis of *C. albicans.* (A) Effects of two different anti-gC1qR monoclonal antibodies on the endocytosis of *C. albicans* by oral epithelial cells. (B) Effects of the anti-gC1qR antibody 74.5.2 and the EGFR kinase inhibitor, gefitinib on the endocytosis of *C. albicans* by oral epithelial cells. (C and D) Endocytosis of *C. albicans* by NIH/3T3 cells expressing human gC1qR and/or human EGFR. (C) Additive effects of EGFR and gC1qR on endocytosis. (D) Effects of inhibiting surface-expressed gC1qR with the anti-gC1qR antibody 74.5.2 on endocytosis. Results are the mean ± SD of three independent experiments, each performed in triplicate. The data were analyzed using one-way analysis of variance with Dunnett’s test for multiple comparisons. ns, not significant; \**P* < 0.05; \*\**P* < 0.01; \*\*\**P* < 0.001.

To investigate the functional relationship between gC1qR and EGFR, we treated the oral epithelial cells with an anti-gC1qR monoclonal antibody, the specific EGFR inhibitor gefitinib, or the antibody and gefitinib in combination. Both the anti-gC1qR antibody and gefitinib inhibited the endocytosis of *C. albicans* similarly, and combining the anti-gC1qR antibody with gefitinib did not decrease endocytosis further (Fig. 3B). None of the treatments altered the number of cell-associated organisms (Fig. S2B). These results suggest that gC1qR may function in the same pathway as EGFR to mediate the endocytosis of *C. albicans*.

We further explored the relationship between gC1qR and EGFR in the endocytosis of *C. albicans* using a heterologous expression approach. We obtained two NIH/3T3 mouse fibroblastoid cell lines, a wild-type cell line and one that had been transfected with human EGFR and HER2 (48). Each of these cell lines was then transfected with either GFP as a control or human gC1qR. When wild-type NIH/3T3 cells were transfected with gC1qR, they endocytosed a similar number of *C. albicans* cells as the control cells (Fig. 3C), indicating that gC1qR was unable to induce endocytosis in the absence of EGFR. As we found previously (13), NIH/3T3 cells that expressed human EGFR and HER2 endocytosed more *C. albicans* cells than the wildtype cells. When the EGFR-expressing cells were transfected with gC1qR, they endocytosed even more organisms. This increase in endocytosis was due to the presence of surface expressed gC1qR because treating these cells with an anti-gC1qR antibody reduced endocytosis to basal levels (Fig. 3D). There was an increase in the number of *C. albicans* cells that were associated with NIH/3T3 cells that expressed EGFR relative to cells that did not (Fig. S2C). However, expression of gC1qR had no significant effect on the number of cell-associated organisms (Fig. S2C, D). Collectively, these data indicate that gC1qR is a key cofactor that enhances EGFR-mediated endocytosis of *C. albicans.*

### Surface-expressed gC1qR is required for *C. albicans-induced* stimulation of epithelial cell production of proinflammatory mediators

To investigate the role of surface-expressed gC1qR in the oral epithelial cell inflammatory response to *C. albicans*, we analyzed the effects of an anti-gC1qR antibody on the secretion of IL-1β and IL-8. In contrast to knockdown of gC1qR with siRNA, inhibiting gC1qR with the monoclonal antibody significantly reduced the secretion of both mediators, indicating that surface expressed gC1qR is required for *C. albicans* to stimulate their production (Fig. 4A and B).

**Fig. 4.**
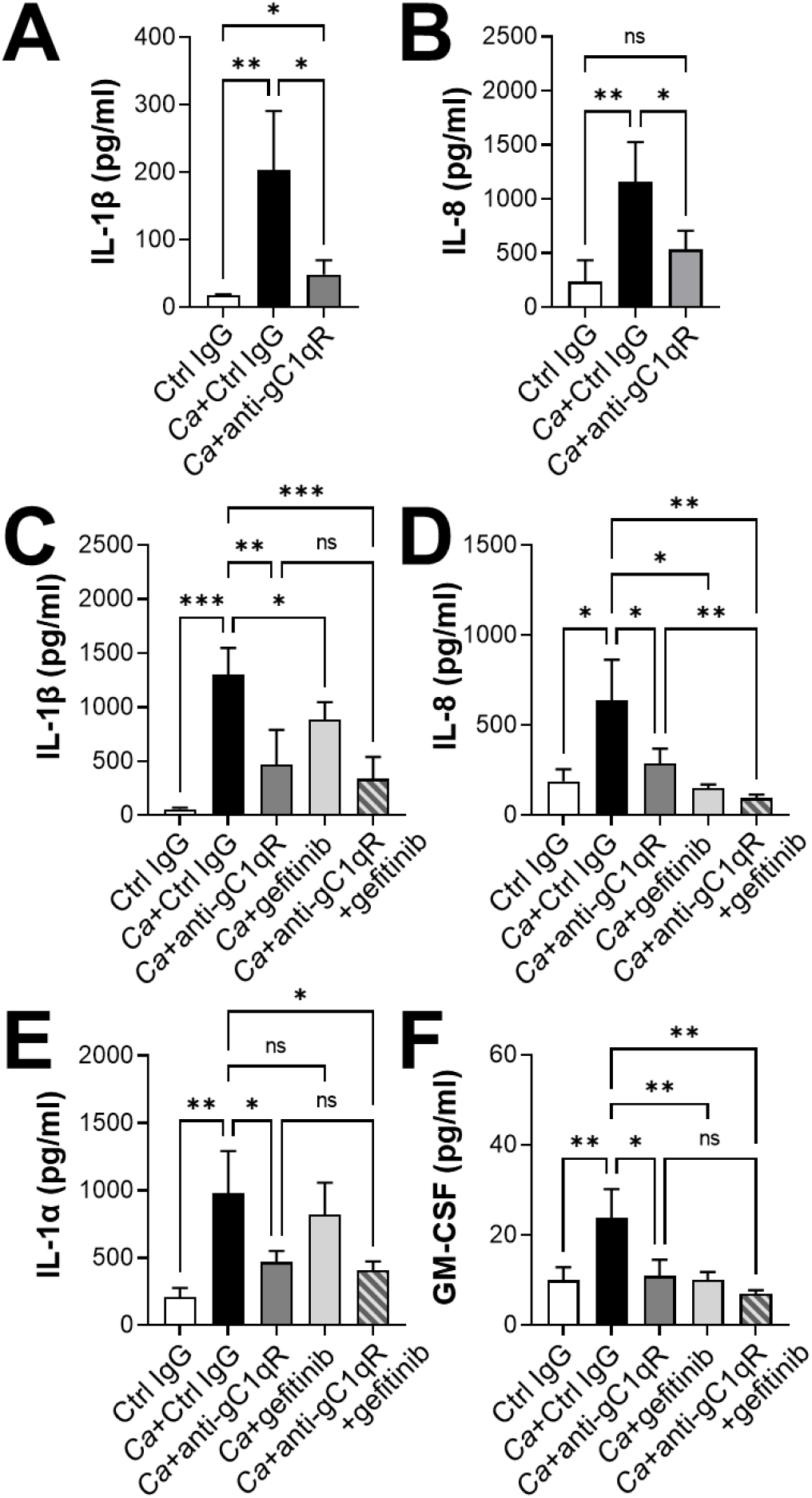
gC1qR is required for production of proinflammatory mediators by oral epithelial cells in response to *C. albicans* infection. (A-F) Oral epithelial cells were infected with *C. albicans* in the presence of an anti-gC1qR antibody 74.5.2 or gefitinib for 8 h, after which the concentration of the indicated inflammatory mediators in the medium was analyzed by ELISA (A and B) or Luminex cytometric bead array (C-F). Results are the mean ± SD of three independent experiments, each performed in duplicate. The data were analyzed using one-way analysis of variance with Dunnett’s test for multiple comparisons. Ca, *C. albicans;* Ctrl, control; ns, not significant; \**P* < 0.05; \*\**P* < 0.01; \*\*\**P* < 0.001.

Next, we analyzed the functional relationship between gC1qR and EGFR in the epithelial cell proinflammatory response. We used a cytometric bead array to measure the levels of IL-1β, IL-8, IL-1α, and GM-CSF that were secreted in response to *C. albicans.* Inhibition of surface-expressed gC1qR significantly reduced the production of all four inflammatory mediators (Fig 4C-F), although the absolute levels of IL-1β and IL-8 were somewhat different when measured by the cytometric bead array instead of the ELISA. Inhibition of EGFR with gefitinib also significantly decreased the levels of IL-1β, IL-8, and GM-CSF, but had no effect on IL-1α levels, as reported previously (15, 16). The finding that inhibition of gC1qR reduced IL-1α production whereas inhibition of EGFR had no effect suggests that gC1qR is required for the production of IL-α by interacting with a receptor other than EGFR.

When the epithelial cells were incubated with the anti-gC1qR antibody and gefitinib in combination, the production of IL-1β and GM-CSF was not inhibited more than in cells incubated with the anti-gC1qR antibody alone (Figs. 4C and F). However, dual inhibition of gC1qR and EGFR resulted in a modest further reduction in IL-8 production (Fig. 4F). Collectively, these results indicate that gC1qR and EGFR function in the same pathway to induce the production of IL-1β, IL-8, and GM-CSF in response to *C. albicans.*

### gC1qR is necessary for intact *C. albicans* to interact with EGFR

Our finding that gC1qR functions in the same pathway as EGFR to mediate epithelial cell endocytosis and stimulation prompted us to investigate whether gC1qR was necessary for *C. albicans* to activate EGFR. We found that treatment of oral epithelial cells with an anti-gC1qR antibody decreased *C. albicans-induced* phosphorylation of EGFR by approximately 58% (Figs. 5A and B). As expected, treatment of the infected cells with the EGFR kinase inhibitor, gefitinib reduced EGFR phosphorylation to below basal levels. Notably, inhibition of gC1qR did not block EGFR phosphorylation in response to either EGF or candidalysin, a pore-forming toxin released by *C. albicans* (15, 49) (Fig. S3). Thus, the effects of inhibiting surface-expressed gC1qR were specific to intact *C. albicans*.

**Fig. 5.**
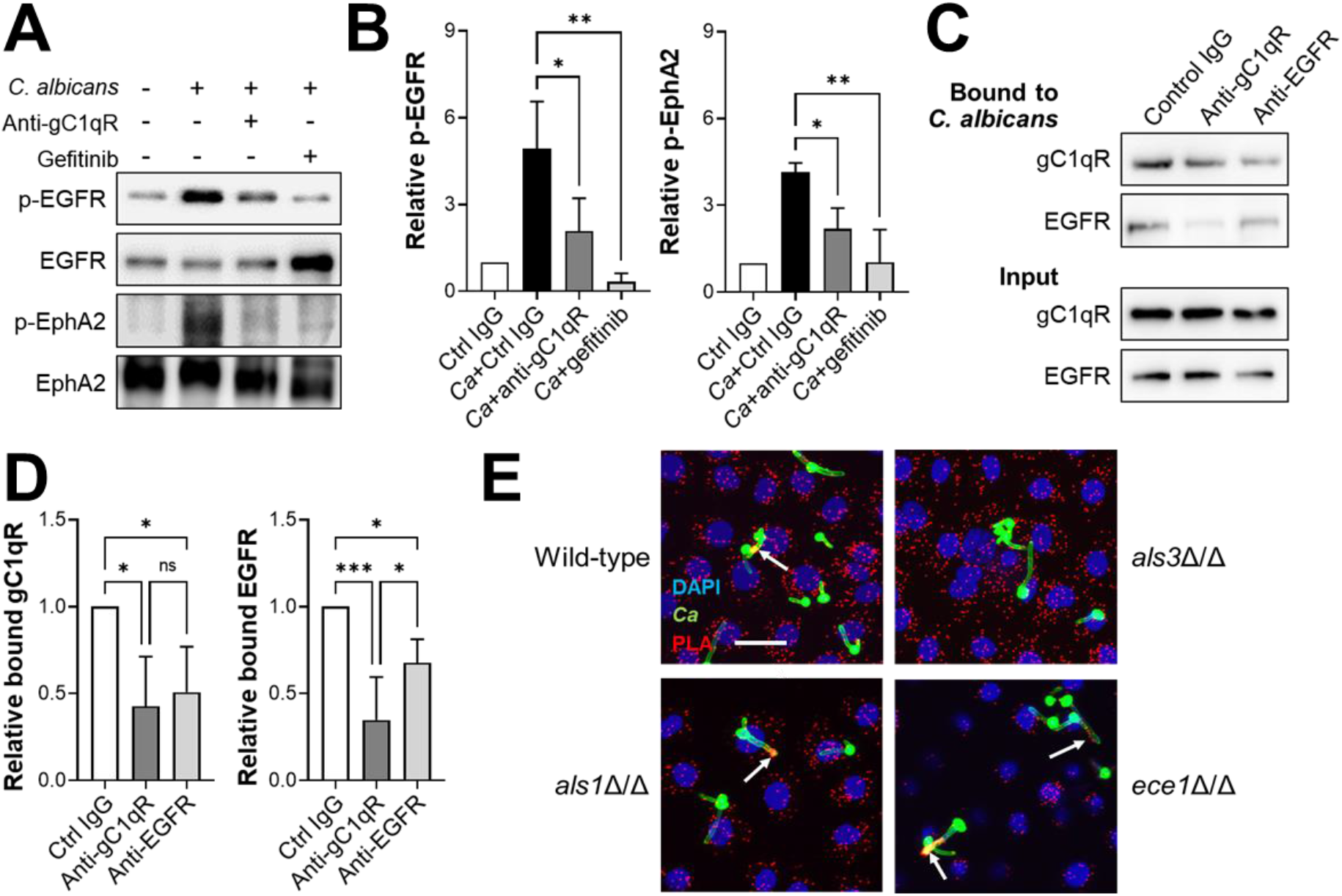
Interactions of gC1qR and EGFR with *C. albicans.* (A and B) Effects of the anti-gC1qR antibody 74.5.2 and gefitinib on the phosphorylation of EGFR and EphA2 in response to 90-min infection with *C. albicans.* (A) Representative immunoblots. (B) Densitometric analysis of three immunoblots, such as the ones shown in (A). (C and D) Effects of anti-gC1qR and anti-EGFR antibodies on binding of gC1qR and EGFR to *C. albicans* hyphae. (C) Representative immunoblots. (D) Densitometric analysis of four immunoblots, such as the ones shown in (C). Results are the mean ± SD of 3-4 independent experiments. (E) Proximity ligation assay showing the association of gC1qR with EGFR around hyphae of *C. albicans* wild-type, *als1*Δ/Δ and *ece1*Δ/Δ strains, but not the *als3*Δ/Δ mutant. Scale bar 25 μm. The numerical data were analyzed using one-way analysis of variance with Dunnett’s test for multiple comparisons. Ca, *C. albicans;* Ctrl, control; ns, not significant; \**P* < 0.05; \*\**P* < 0.01; \*\*\**P* < 0.001.

Activation of EGFR is also required for *C. albicans* to induce sustained phosphorylation of the EphA2 receptor tyrosine kinase (11, 16). We determined that inhibition of gC1qR decreased *C. albicans*-induced EphA2 phosphorylation (Figs. 5A and B). These findings are consistent with the model that gC1qR is necessary for *C. albicans* to activate EGFR and its downstream signaling pathways in oral epithelial cells.

Previously, we found that EGFR associated, either directly or indirectly, with *C. albicans* hyphae (13). To determine if gC1qR plays a role in this association, we infected oral epithelial cells with *C. albicans* in the presence of either an anti-gC1qR antibody or an anti-EGFR antibody. After 90 min, we lysed the epithelial cells with a detergent, collected the *C. albicans* hyphae, and rinsed them extensively to remove unbound proteins. Using high molar urea, we eluted the epithelial cell proteins that remained associated with the organisms and analyzed them by immunoblotting. In control cells, both gC1qR and EGFR were associated with the *C. albicans* hyphae (Figs 5C and D). When the cells were incubated with the anti-gC1qR antibody, there was a significant reduction in the amounts of gC1qR and EGFR that were associated with *C. albicans.* When the cells were incubated with an anti-EGFR antibody, the amount of fungal-associated gC1qR and EGFR was also significantly reduced. Collectively, these data suggest that gC1qR and EGFR have a reciprocal relationship such that each protein is necessary for the other to maximally associate with *C. albicans*.

### *C. albicans* Als3 but not candidalysin is required for increased association of gC1qR with EGFR

Both the Als3 invasin and the candidalysin pore forming toxin are required for *C. albicans* to maximally activate EGFR in oral epithelial cells (15, 16). We investigated whether these factors were required gC1qR to associate with EGFR around *C. albicans* cells in intact oral epithelial cells. Using the proximity ligation assay, we determined that when oral epithelial cells were infected with an *als3*Δ/Δ deletion mutant, there was no accumulation of the gC1qR-EGFR containing complex around the hyphae (Fig. 5E). By contrast, when the epithelial cells were infected with the wild-type strain, an *als1Δ/Δ* deletion mutant, or a candidalysin-deficient *ece1Δ/Δ* deletion mutant, this complex formed around the hyphae. Thus, Als3, but not Als1 or candidalysin, is necessary for gC1qR to associate with EGFR around *C. albicans* hyphae.

### Inhibition of gC1qR does not alter *C. albicans* virulence or activation of EGFR in mice

Next, we investigated the role of gC1qR in the pathogenesis of oropharyngeal candidiasis in mice. To verify that monoclonal antibodies raised against human gC1qR could inhibit mouse gC1qR, we tested the capacity of these antibodies to inhibit the endocytosis of *C. albicans* by primary mouse epithelial cells. We found monoclonal antibody 74.5.2, which is directed against the binding site for high-molecular weight kininogen (47) significantly reduced *C. albicans* endocytosis (Fig. 6A). It also decreased the number of cell-associated organisms (Fig. S4). By contrast, monoclonal antibody 60.11, which recognizes the binding site for C1q (47), had no detectable effect on either interaction.

**Fig. 6.**
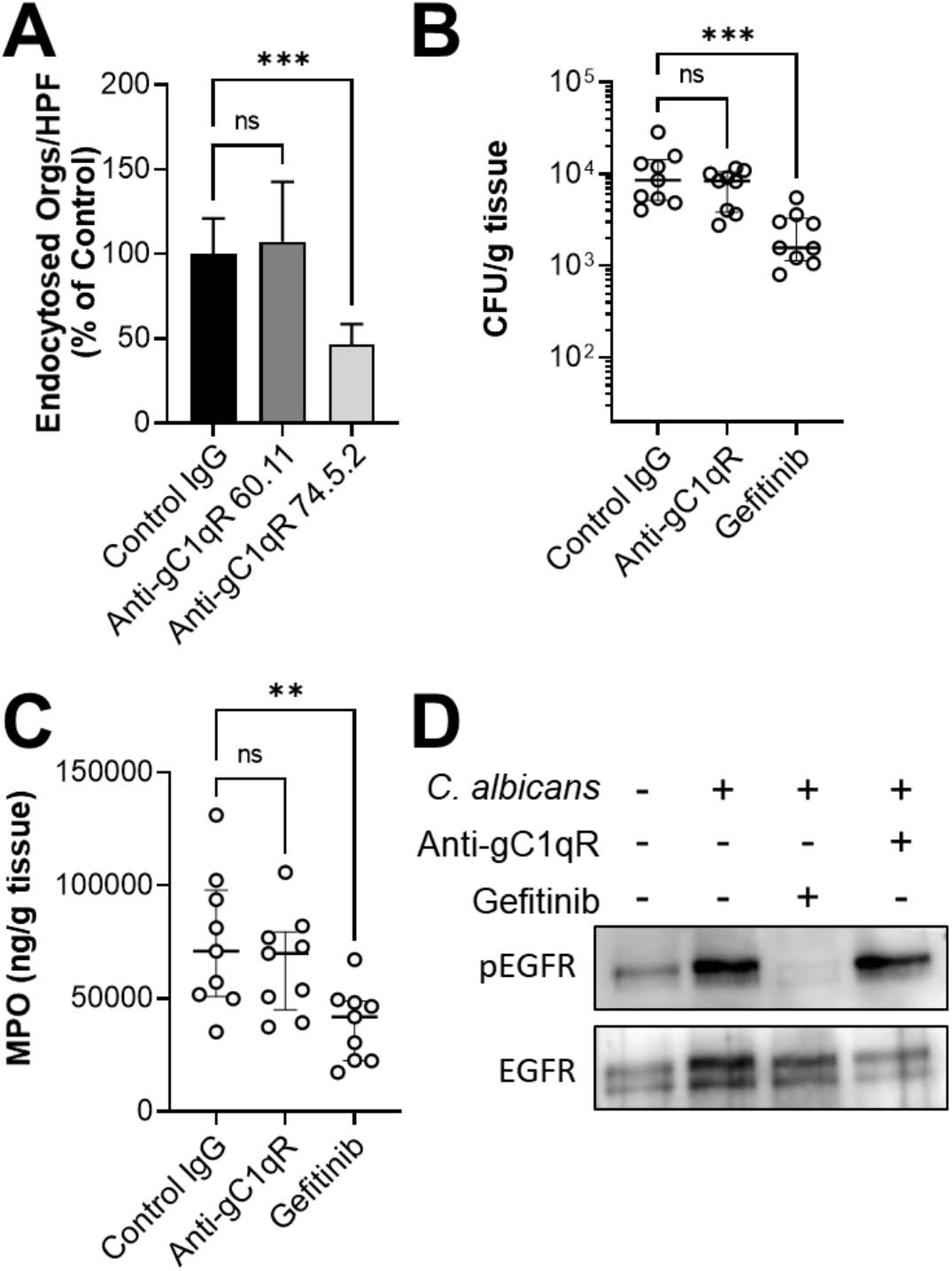
Inhibition of gC1qR has no significant effect on the outcome of oropharyngeal candidiasis. (A) Effects of the indicated anti-gC1qR antibodies on the endocytosis of *C. albicans* by primary mouse oral epithelial cells. Results are the mean ± SD of three independent experiments, each performed in triplicate. The data were analyzed using one-way analysis of variance with Dunnett’s test for multiple comparisons. (B and C) Effects of the anti-gC1qR antibody 74.5.2 and gefitinib on the outcome of oropharyngeal candidiasis after 1 d of infection. Oral fungal burden (B). Oral myeloperoxidase (MPO) content (C). Results are the median ± interquartile range of two independent experiments, each with 4-5 mice per group. The data were analyzed using the Kruskal-Wallis test. (D). Immunoblot showing that the anti-gC1qR antibody 74.5.2 does not block *C. albicans-induced* phosphorylation of EGFR in primary mouse oral epithelial cells. Results are representative of three independent experiments. ns, not significant; \*\**P* < 0.01; \*\*\**P* < 0.001.

Based on these results, we treated immunocompetent mice with antibody 74.5.2 and orally infected them with wild-type *C. albicans.* Control mice were treated with either mouse IgG or gefitinib. We found that after 1 day of infection, mice treated with the anti-gC1qR antibody had the same oral fungal burden as mice that received the control IgG (Fig. 6B). The level of myeloperoxidase (MPO), a measure of phagocyte accumulation (50, 51), in the oral tissues was also not changed by administration of the anti-gC1qR antibody (Fig. 6C). As expected (16), treatment with gefitinib significantly reduced both the oral fungal burden and oral MPO levels in the mice. These results suggest the gC1qR is dispensable for mediating the epithelial cell response to *C. albicans* in mice.

To investigate these results further, we analyzed the capacity of the anti-gC1qR antibody 74.5.2 to inhibit *C. albicans-induced* phosphorylation of EGFR in mouse epithelial cells. We determined that the antibody had no effect, although EGFR phosphorylation was inhibited by gefitinib (Fig. 6D). Thus in mice, gC1qR is likely dispensable for *C. albicans-induced* EGFR activation, unlike in humans.

## DISCUSSION

In this work, we determined that EGFR interacts with numerous proteins that function in a multitude of different signaling pathways, consistent with the concept that this receptor is a key regulator of epithelial cell physiology. The proteomics data indicate that EGFR is part of a multi-protein complex that contains E-cadherin, EphA2, and HERS. This result provides an explanation for our previous findings that these three receptors function with EGFR in the same pathway to induce the epithelial cell response to *C. albicans* (11, 13, 18).

Previously, we had determined that *C. albicans* activates the aryl hydrocarbon receptor, leading to the de-repression of src family kinases that phosphorylate EGFR (14). However, the member(s) of the src family responsible for this phosphorylation remained unknown. The current finding indicate that Lyn and Yes associate with EGFR suggests that these two members of the src family kinases phosphorylate EGFR in response to *C. albicans* infection.

EGFR mediates the endocytosis of *C. albicans* by activating the cortactin-clathrin pathway, leading to the formation of pseudopods that engulf the organism and pull it into the epithelial cell (21, 22). The proteomics data were consistent with this mechanism and suggest that the arp2/3 complex mediates the rearrangement of actin filaments that induce pseudopod formation. Although the arp2/3 complex can be activated by the Wiskott-Aldrich syndrome protein (WASP) or vasodilator-stimulated phosphoprotein (VASP) (52–54), neither protein was consistently associated with EGFR in cells infected with *C. albicans.* Thus, it remains to be determined how EGFR activates this complex

We also established roles for four key proteins that interact with EGFR and induce the epithelial cell response to *C. albicans,* none of which had been previously implicated in the response to fungal pathogens. Three of these proteins, WBP2, IFITM3, and gC1qR, were required for maximal epithelial cell endocytosis of the fungus, suggesting that they function along with EGFR to orchestrate this process.

Perhaps not surprisingly, siRNA knockdown of these proteins had pleotropic effects on the production of IL-1β and IL-8 by the infected oral epithelial cells, as these are regulated by a myriad of inflammatory and infectious signals. Knockdown of WBP2 led to reduced cellular levels of EGFR and thus resulted in decreased production of IL-1β and IL-8, similar to siRNA knockdown of EGFR itself. This result is consistent with a report that WBP2 is required for normal EGFR expression (32). Knockdown of TOLLIP inhibited the production of IL-8 but had no effect on IL-1β secretion. Although TOLLIP is generally considered to be a negative regulator of the host inflammatory response (35–37), our results suggest that it has the capacity to be a positive regulator of IL-8 production in epithelial cells infected with *C. albicans.* While siRNA knockdown of IFITM3 inhibited *C. albicans* endocytosis, it stimulated the production of both IL-1β and IL-8. IFITM3 has been found to enhance the degradation of activated EGFR in pulmonary epithelial cells (38). Although knockdown of IFITM3 in oral epithelial cells had no detectable effect on total cellular EGFR levels, our results indicate that IFITM3 is a positive regulator of epithelial cell endocytosis of *C. albicans* but a negative regulator of cytokine production.

Our in-depth analysis of gC1qR enabled us to develop a model for how this protein interacts with EGFR and is required for EGFR signaling in response to *C. albicans* infection (Fig. 7). We propose that gC1qR and EGFR form part of a complex that interacts either directly or indirectly with the *C. albicans* Als3 invasin. According to this model, gC1qR is required for Als3 to maximally interact with and activate EGFR. In turn, EGFR is required for *C. albicans* to interact optimally with gC1qR. This arrangement is deduced from the proteomics analysis, which showed that gC1qR was associated with EGFR, and from pull-down assays, which indicated that blocking gC1qR inhibited the physical association of EGFR with *C. albicans* and that blocking EGFR in turn reduced the association of gC1qR with the organism. Although the proteomics analysis suggested that gC1qR associated with EGFR constitutively, the proximity ligation assay showed that the gC1qR-EGFR complex was enriched in regions surrounding *C. albicans* hyphae that expressed Als3. Previously, we found that Als3 is required for *C. albicans* to induce maximal EGFR activation (13). Accordingly, these results suggest that gC1qR is a key co-factor that is required for Als3 to activate EGFR (Fig. 7).

**Fig. 7.**
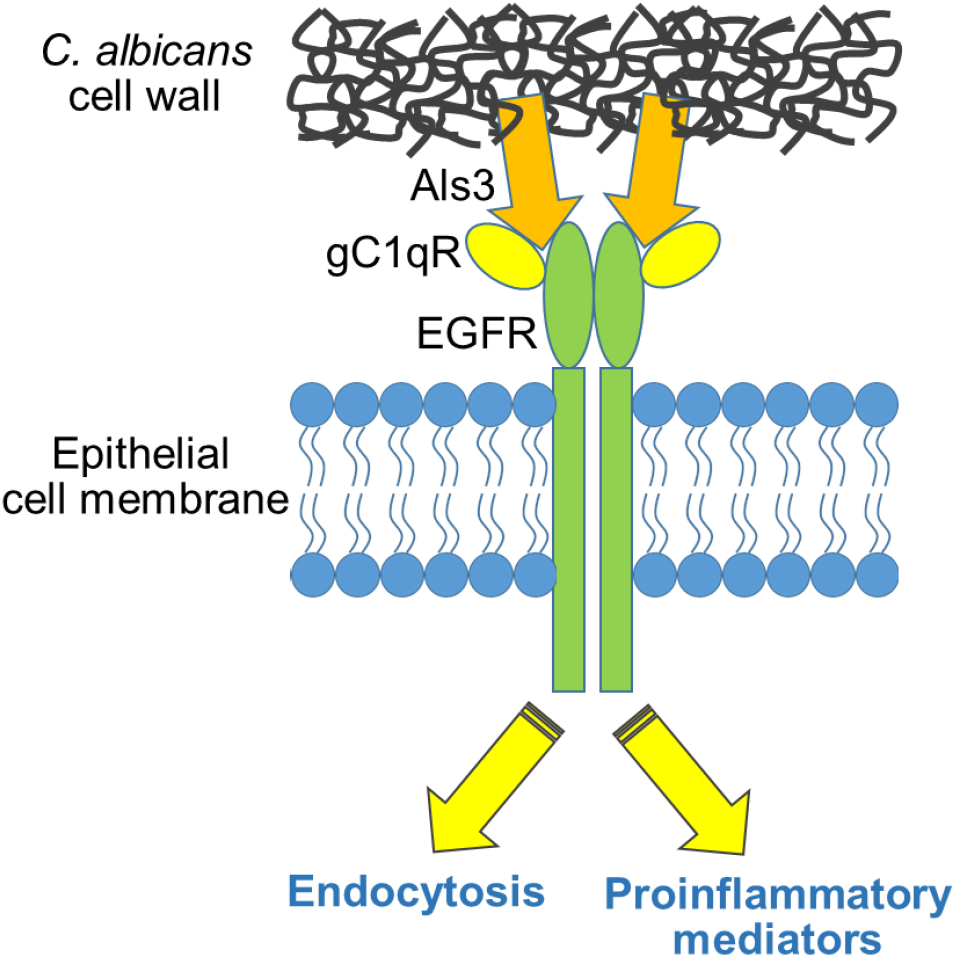
Diagram of the functional interaction of *C. albicans* Als3 with gC1qR and EGFR in human oral epithelial cells. Als3 interacts either directly or indirectly with both gC1qR and EGFR, leading to the activation of EGFR and subsequent induction of endocytosis and secretion of proinflammatory mediators.

Although EGF also activates EGFR, we found that it does so independently of gC1qR. These findings differ from Kim et al. (46), who reported that inhibition of gC1qR in the A549 lung cancer cell line blocked activation of EGFR induced by EGF. Possible reasons for these discrepant results include differences in the cell lines and the anti-gC1qR antibodies that we used as compared those used by the other investigators. We also found that blocking gC1qR did not prevent activation of EGFR by candidalysin. Collectively, these data suggest that soluble stimuli activate EGFR in oral epithelial cells independently of gC1qR whereas *C. albicans* hyphae that express Als3 activate EGFR by a mechanism that requires gC1qR.

When investigating the role of gC1qR in mediating the production of IL-1β and IL-8 in response to *C. albicans* infection, we observed that treatment with the anti-gC1qR antibody inhibited the production of these inflammatory mediators whereas siRNA knockdown of gC1qR had the opposite effect. The likely explanation for these results is that inhibiting just extracellular gC1qR has a different effect than decreasing the levels of intracellular gC1qR. Our experiments with the anti-gC1qR antibody indicate that it inhibits the production of proinflammatory mediators by blocking *C. albicans*-induced activation of EGFR and EphA2. These latter two receptors play key roles in stimulating the production of cytokines and chemokines in response to *C. albicans* (15, 16). Why reducing the levels of intracellular gC1qR stimulates the production of proinflammatory mediators in oral epithelial cells is unknown. However, knockdown of gC1qR in macrophages is known to reduce the cytoplasmic levels of tumor necrosis factor α inducible protein 3 (TNFAIP3, A20), a suppressor of NF-κB activation, thereby stimulating the production of proinflammatory mediators (55). We speculate that a similar process may occur when gC1qR is knocked down in oral epithelial cells.

Inhibition of EGFR in mice with oropharyngeal candidiasis reduces oral fungal burden and inflammation (16). Thus, we were surprised to determine that treating mice with the anti-gC1qR antibody had no effect on either of these parameters. Subsequently, we found that this antibody did not block *C. albicans*-induced phosphorylation of EGFR in primary oral epithelial cells. The probable explanation for these results is that gC1qR is dispensable for the fungus to activate EGFR in mouse epithelial cells. A less likely possibility is that the anti-gC1qR antibody used in the mouse studies did not block the association of gC1qR with EGFR in mouse cells, even though it did so in human cells.

gC1qR is exploited by multiple microbial pathogens. gC1qR mediates the adherence of *Plasmodium falciparum* and *Staphylococcus aureus* to vascular endothelial cells and the adherence of *Bacillus cereus* spores to epithelial cells (44, 45, 56, 57). It is also an epithelial cell receptor for the *L. monocytogenes* internalin B (InlB) invasin and mediates the endocytosis of the organism (28). Because gC1qR lacks a transmembrane sequence, it is thought to induce endocytosis by signaling via another cell surface receptor. In the case of *L. monocytogenes,* the second receptor appears to be Met, the hepatocyte growth factor receptor (58). In some cell lines, gC1qR and Met have been found to function cooperatively to mediate the endocytosis of this bacterium (59). Although the primary amino acid sequence of the *C. albicans* Als3 invasin shares no homology with InlB, we demonstrate that Als3 also interacts with gC1qR. Instead of Met, EGFR is the second receptor that associates with gC1qR and transduces the epithelial cell response *C. albicans.* Not only is gC1qR required for EGFR-mediated endocytosis of the fungus, but it is also necessary for the maximal epithelial cell inflammatory response to this organism. We have identified additional host cell receptors for *C. albicans* including E-cadherin, HER2, the platelet-derived growth factor BB, the aryl hydrocarbon receptor, and gp96 (12–14, 18, 60). Whether gC1qR also interacts with these receptors remains to be determined.

## MATERIALS AND METHODS

### Epithelial cells and fungal strains

The OKF6/TERT-2 oral epithelial cell line was provided by J. Rheinwald (Dana-Farber/Harvard Cancer Center, Boston, MA) and cultured as outlined previously (14, 17). Primary oral mucosal epithelial cells from BALB/c mice were obtained from Cell Biologics Inc. and grown following the manufacturer’s instructions. NIH/3T3 cells that expressed human EGFR and HER2 were provided by Nadege Gaborit (Institut de Researche en Cancérologie de Montpellier, France) and grown as described (48). The *C. albicans* wild-type strain SC5314 and the *als1Δ/Δ, als3Δ/Δ,* and *ece*1Δ/Δ mutants (16) were grown in yeast extract, peptone, dextrose (YPD) broth in a shaking incubator at 30°C for 18 h, after which the cells were pelleted by centrifugation and washed twice with PBS. Yeast cells were suspended in PBS, diluted and counted with a hemacytometer.

### Immunoprecipitation

OKF6/TERT-2 oral epithelial cells were grown to confluency in 75 cm^2^ tissue culture flasks and the culture medium was changed to KSF medium without supplements (Thermo Fisher Scientific; # 17005042) the night before the experiment. The next morning, cells were incubated for 90 min with supplement-free KSF medium containing 10^8^ *C. albicans* yeast cells or for 5 min with 50 ng/ml EGF. Cell incubated with fresh supplement-free KSF alone were processed in parallel as a negative control. After incubation, the medium was aspirated and the cells were washed once with ice cold PBS with Ca^2+^ and Mg^2+^ (PBS^++^). The proteins were cross-linked by incubation with 1% paraformaldehyde for 10 min. After rinsing the cells twice with ice cold PBS^++^, they were detached from the culture flasks with a cell scraper and collected by centrifugation. The cell pellet was lysed with 5.8% octyl-β-D-glucopyranoside (VWR; # 97061-760) with sonication. The lysate was clarified by centrifugation and then precleared with protein A/G magnetic beads (Thermo Fisher Scientific, # PI88802,) for 30 min at 4°C. The precleared cell lysates were incubated with an anti-EGFR antibody (Santa Cruz Biotechnology; # SC-101, clone R-1), for 1 h at 4°C, and precipitated with protein A/G magnetic beads for 2 h at 4°C. After washing the beads 3 times with 1.5% octyl-β –D-glucopyranoside, the proteins were eluted with 8M Urea, stored at −80°C.

### Mass Spectrometry

The protein samples were treated with Tris (2-carboxyethyl) phosphine hydrochloride (Thermo Fisher Scientific; # 20491) and 2-chloroacetamide (cat# 154955) for reduction and alkylation, respectively. Tris buffer (0.1 M, pH 8.5) was added to each sample decrease the urea concentration to final 2 M. After dilution, the samples were digested with lysyl endopeptidase (Fujifilm Wako Chemicals; # 125-05061) and trypsin (Thermo Fisher Scientific; # 90058) for 20 h at 37°C. After quenching the reaction by adjusting the pH to 3, the digested samples were desalted using C18 columns (Thermo Fisher Scientific; # 89870) and then dried by vacuum centrifuge.

For the LC-MS/MS analysis, the samples were dissolved in 0.2% formic acid (FA) solution. The analysis was performed using an EASY-nLC 1000 (Thermo Fisher Scientific connected to a Q Exactive Orbitrap Mass Spectrometers (Thermo Fisher Scientific). Solvent A consisted of 99.9% H2O and 0.1% FA, and solvent B consisted of 19.9% H2O, 80% acetylnitrile, and 0.1% FA. Each sample was loaded onto an Easy Spray Column (25 cm × 75 μm, 2 μm C18, ES802, Thermo Fisher Scientific) and separated over 90 min at a flow rate of 0.5 μL/min with the following gradient: 2-35% B (75 min), 35-85% B (5 min), and 85% B (10 min). The full MS scan was acquired at 70,000 resolution with a scan range of 350-2000 m/z, the automatic gain control target was 1 × 10^6^, and the maximum injection time was 100 ms. The dd-MS2 scan was acquired at 17,500 resolution with a scan range of 200-2000 m/z, the automatic gain control target was 5 × 10^4^, the maximum injection time was 64 ms, and the isolation window was 2.0 m/z. The proteomic data processing was performed using Proteome Discoverer 1.4 (Thermo Fisher Scientific) and the Sequest HT Search Engine. The search allowed for a precursor mass tolerance of 10 ppm, a minimum peptide length of 6, and a minimum peptide sequence number of 1.

### Proximity ligation assay

To determine if the selected proteins associated with EGFR in intact epithelial cells, OKF6/TERT-2 cells were grown to confluency on fibronectin coated coverslips. The cells were infected with 3×10^5^ *C. albicans* yeast in supplement-free KSF medium. After 90 min, the medium was aspirated and the cells were fixed with 4% paraformaldehyde for 10 min. The coverslips were washed 3 times with PBS, after which the epithelial cells were permeabilized with 0.1% triton X-100 in PBS for 20 min. The association between EGFR and WBP2, GBP6, TOLLIP, EEA1, DSG3, IFITM3, or gC1qR, was detected using the Duolink in Situ Red Starter Kit Mouse/Rabbit (Sigma-Aldrich; #DUO92101-1kit) according to the manufacturer’s instruction. The antibodies used were rabbit anti-EGFR (Genetex; # GTX121919, clone N1-2), mouse anti-EGFR (Santa Cruz Biotechnology # SC-101, clone R-1), anti-WBP2 (Santa Cruz Biotechnology; # SC-514247, clone D-12), anti-GBP6 (Sigma-Aldrich; # HPA027744), anti-TOLLIP (Proteintech # 117315-1), anti-EEA1 (Santa Cruz Biotechnology; # SC-365652 clone E-8), anti-DSG3 (Santa Cruz Biotechnology; # SC-53487, clone 5g11), anti-IFITM3 (Proteintech; # 11714-1), and gC1qR (Santa Cruz Biotechnology; # SC-23885, clone 74.5.2). The cells were imaged by confocal microscopy and z stacks of the images were constructed using the Leica Application Suite X software (Leica).

### siRNA

To knock down the levels of selected proteins, the OKF6/TERT-2 cells were grown in 6-well tissue culture plates to 50-80% confluency overnight. The next morning, the cells were transfected with EGFR (Santa Cruz Biotechnology; # SC-29301), WBP2 (Santa Cruz Biotechnology; # SC93955), TOLLIP (Dharmacon; # L-016930-00-0005), IFITM3 (Santa Cruz Biotechnology; # SC-97053), gC1qR (Santa Cruz Biotechnology; # SC-42880), or control (Qiagen; # 1027281) siRNA using Lipofectamine RNAiMAX (Invitrogen; #13778150) following the manufacturer’s instructions. After 24 h post-transfection, the cells were seeded onto fibronectin coated coverslips and incubated for another 24 h before use in the experiments. The extent of protein knockdown was determined by immunoblotting and total loading was detected by probing the blots with an anti-GAPDH antibody (Cell Signaling; #5174, clone D16H11).

### Epithelial cell endocytosis and cell-association

Our standard differential fluorescence assay was used to measure the number of *C. albicans* cells that were endocytosed by and cell-associated with the OKF6/TERT-2 epithelial cells, primary mouse epithelial cells, and NIH/3T3 cells (13, 18, 22). The host cells were infected with 10^5^ *C. albicans* yeast cells. To ensure similar levels of endocytosis among the different host cells, the incubation time was 2.5 h for OKF6/TERT-2 cells, 2 h for the mouse oral epithelial cells and 1.5 h for the NIH/3T3 cells. In the siRNA experiments, the transfected epithelial cells were seeded onto fibronectin-coated glass coverslips 24 h before infection. For experiments using gefitinib (1 μM; Selleck Chem, Inc; # S1025) or the anti-gC1qR antibodies (10 μg/ml), the inhibitor or antibodies were added to the host cells 1 h prior to infection and they remained in the medium for the duration of the experiment. All experiments performed in triplicate at least three times.

### Lentivirus construction and production and host cell transduction

The transfer vectors (pLenti-EF1A-EGFP-Blast or pLenti-EF1A-hC1QBP[NM_001212.4]-Blast) were constructed by cloning eGFP or hC1QBP[NM_001212.4] into pLenti-Cas9-Blast (Addgene; # 52962) at the BamHI and XbaI sites. The virus was produced by transfecting HEK293T cells with plasmid psPAX2 (Addgene; # 12260), plasmid pCMV-VSVG (Addgene; # 8454), and transfer vector (pLenti-EF1A-EGFP-Blast or pLenti-EF1A-hC1QBP[NM_001212.4]-Blast) using the X-tremeGENE 9 DNA transfection reagent (Sigma-Aldrich; # 6365787001) according to the manufacturer’s instructions. The supernatant containing the virus was collected at 60 h posttransfection, passed through a 0.45um PVDF filter and stored at 4°C (short-term) or −80°C (longterm).

For transduction, the NIH/3T3 mouse fibroblast cells (untransformed control cells or the human EGFR/hHER2 transformed cell line (48)) were seeded into a 6-well plate in DMEM+10% bovine calf serum. The cells were transduced with lentivirus in the presence of 0.5μg/ml polybrene (Santa Cruz Biotechnology; #SC134220). The plates were centrifuged at 1000*g* for 30 min and then incubated at 37°C in 5%CO2 overnight. The next morning, the cells were transferred to 10 cm diameter tissue culture dishes. Two days post transduction, 10 μg/ml of blasticidin (Gibco; # A1113903) was added to the medium to select for transduced cells and selection was maintained for 7 days. The successful transduction of eGFP was determined by fluorescent microscopy and transduction of hC1QBP (gC1qR) was verified via immunoblotting with an anti-gC1qR antibody (clone 74.5.2) and an anti-EGFR antibody (Cell Signaling Technology; # 4267, clone 38B1).

### Receptor phosphorylation

Analysis of the phosphorylation of EGFR and EphA2 was performed as previously described (11, 16). Briefly, OKF6/TERT-2 cells were seeded onto 24-well tissue culture plates and incubated overnight in supplement-free KSF medium. The next morning, the cells were treated with gefitinib, an anti-gC1qR antibody (clone 74.5.2) or control mouse IgG (R&D systems; #MAB002) for 1 h. The cells were then stimulated with 10^6^ *C. albicans* yeast cells for 90 min, 40 μM candidalysin (Biomatik) for 5 min, or 1 ng/ml EGF for 5 min in the presence of the inhibitor or antibody. Next, the cells were lysed with 100μl 2X SDS loading buffer in the present of phosphatase inhibitor cocktail (Sigma-Aldrich), protease inhibitor cocktail (Sigma-Aldrich), and PMSF (Sigma-Aldrich). After the samples were denatured at 90°C for 2 min, they were clarified by centrifugation. The proteins were separated by SDS-PAGE and transferred to PVDF membranes. Phosphorylated EGFR (Tyr 1068) was detected with a phosphospecific antibody (Cell Signaling Technology; # 2234) and enhanced chemiluminescence. Total EGFR was detected with an anti-EGFR antibody Cell Signaling Technology; # 4267). Phosphorylation of EphA2 (Ser 897) was detected with a phosphospecific anti-EphA2 antibody (Cell Signaling Technology; #6347, clone D9A1) and total EphA2 was detected with an anti-EphA2 antibody (Cell Signaling Technology; # 6997, clone D4A2). The phosphorylation of EGFR in mouse oral epithelial cells was determined similarly except that total EGFR was detected with anti-EGFR antibody from Santa Cruz Biotechnology (# SC373746, clone A-10). All experiments were repeated at least three times.

### *C. albicans* association with epithelial cell gC1qR and EGFR

The capacity of anti-gC1qR and anti-EGFR antibodies to block the association of *C. albicans* with gC1qR and EGFR was determined using our previously described method (61). Confluent OKF6/TERT-2 epithelial cells in a 6-well tissue culture plate were switched to supplement-free KSF medium the night before the experiment. The next morning, the epithelial cells were incubated with 10 μg/ml of control mouse IgG, an anti-gC1qR antibody (clone 74.5.2), or an anti-EGFR antibody (cetuximab; Lilly) for 1 h, and then infected with 6×10^6^ *C. albicans* blastospores. After 90 min of infection, the cells were detached with a cell scraper and collected by centrifugation. The epithelial cells were lysed by incubation with 5.8% n-octyl-β glucopyranoside in PBS^++^ in the present of protease inhibitors and PMSF on ice for 1 h. The samples were centrifuged at 5,000 g for 1 min and the supernatants containing the total cell lysates were collected. The pellets, which contained the intact *C. albicans* cells and the epithelial cells proteins that were associated with them, were washed three times with cold 1.5% n-octyl-β D glucopyranoside in PBS^++^ containing protease inhibitors. The proteins that remained associated with the *C. albicans* cells were eluted by incubation with 6M urea for 30 min on ice. After pelleting the organisms by centrifugation, the eluted proteins in the supernatants were separated by SDS PAGE. The presence of gC1qR and EGFR in the eluates was determined by immunoblotting with an anti gC1qR antibody (Abcam; # ab24733, clone 60.11) and an anti EGFR antibody (clone 38B1). The experiment was repeated four times.

### Cytokine and chemokine measurement

To measure the production of IL-1β and IL-8 by oral epithelial cells, the OKF6/TERT-2 cells were grown in 48-well tissue culture plates overnight in supplement-free KSF and then infected with 6.25 × 10^5^ *C. albicans* yeast cells for 8 h. The conditioned medium was collected and clarified by centrifugation. The levels of IL-1β and IL-8 in the conditioned medium were measured by ELISA using the human IL-1β DuoSet (R&D Systems; # DY20105) and the human IL-8 set (BD Biosciences; # BD555244).

The production of IL-1β, IL-8, IL-1α, and GM-CSF by the epithelial cells was measured similarly except that the epithelial cells were grown in 24-well tissue culture plates and infected with 1.5×10^6^ *C. albicans* yeast cells. The levels of the inflammatory mediators were measured by Luminex Multiplex (R&D systems; # LXSAHM-06). The experiments were repeated three times in duplicate.

### Mouse experiments

Our standard immunocompetent mouse model of oropharyngeal candidiasis was used to assess the effects of an anti-gC1qR antibody and gefitinib the infection (11, 16, 62). Immunocompetent Balb/c were administered 100 μg of the anti-gC1qR antibody (clone 74.5.2) by intraperitoneal injection on day −1 relative to infection. Control mice were injected with a similar amount of mouse IgG. Gefitinib was administered by adding the drug to powdered chow to a final concentration of 200 parts per million, starting at day −2 relative to infection and continuing throughout the experiment. On the day of the infection, the mice were sedated and a calcium alginate swab saturated with 2 × 10^7^ *C. albicans* yeast cells was placed sublingually for 75 min. After 1 day of infection, the mice were sacrificed and the tongues were excised, weighed, and homogenized. The oral fungal burden was determined by quantitative culture of an aliquot of the homogenates and the level of MPO in the homogenates was determined by commercial ELISA (Hycult Biotech; # HK210-02). The experiment was repeated twice using 4-5 mice per condition and the results were combined.

### Statistical analysis

The *in vitro* data were analyzed using one-way analysis of variance with Dunnett’s test for multiple comparisons. The mouse data were analyzed by the Kruskal-Wallis test with Dunn’s test for multiple comparisons. *P* values < 0.05 were considered to be significant.

## ACKNOWLEGEMENTS

We thank Adam Diab for assistance with tissue culture. The work was support in part by NIH grants R01DE026600 and R01AI124566 to SGF, R00DE026856 to MS, and DE022550 to SLG. The QE Orbitrap used in these experiments was purchased with funds from the Department of Pediatrics at Harbor-UCLA Medical Center and the Henry L. Guenther Foundation.

---

**Fig. S1.**
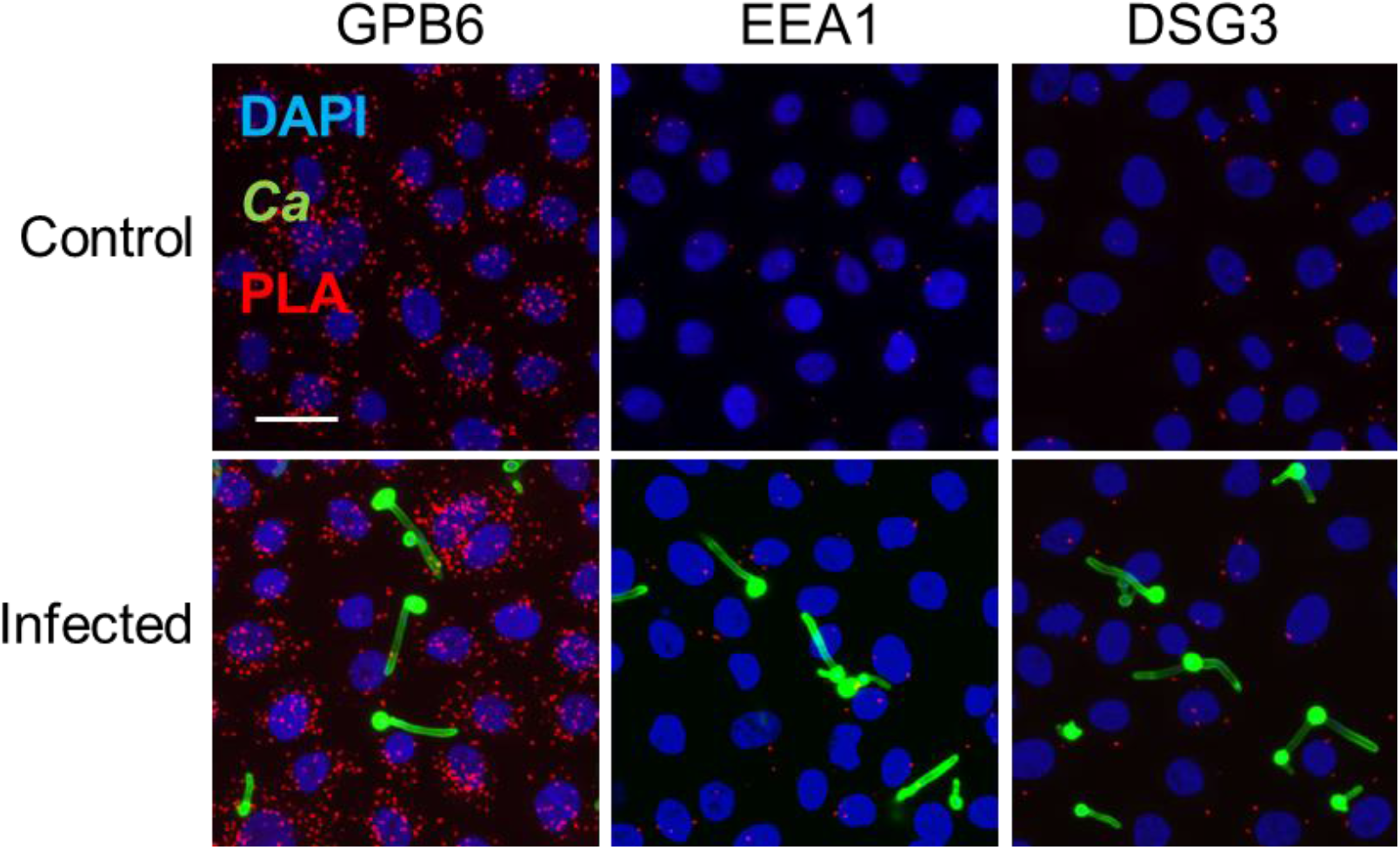
Proximity ligation assays to assess the physical association of the epidermal growth factor receptor (EGFR) with guanylate binding protein 6 (GBP6), early endosome antigen 1 (EEA1), and desmoglein-3 (DSG3) in the OKF6/TERT-2 oral epithelial cell line. The epithelial cells were incubated in either medium alone (top) or infected with *C. albicans* (bottom) for 90 min. Red spots indicate the regions where the indicated proteins associate with EGFR. Results are representative of three independent experiments. Scale bar 25 μm.

**Fig. S2.**
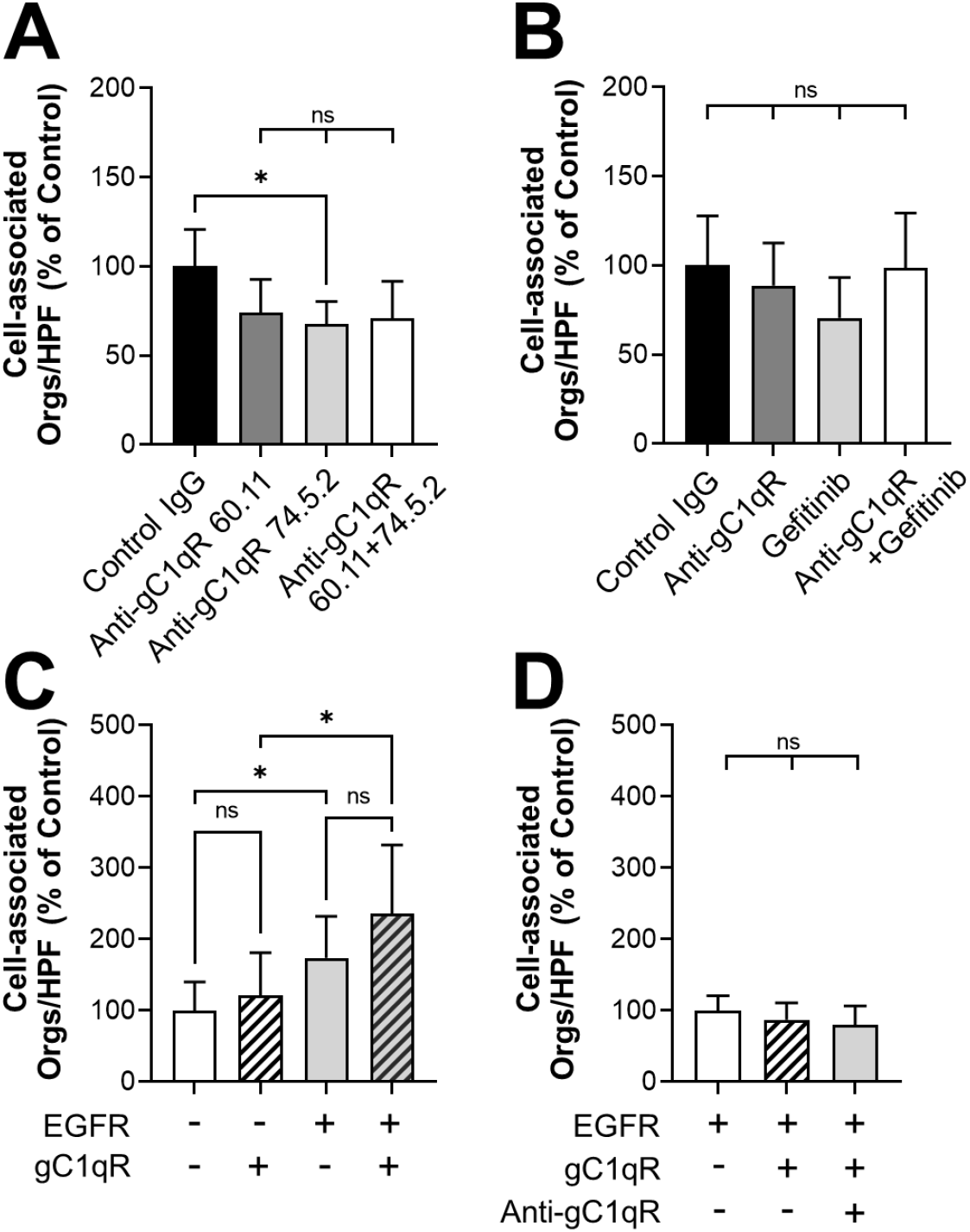
Surface-expressed gC1qR has minimal effects on the number of *C. albicans* cells that are associated with oral epithelial cells. (A) Effects of two different anti-gC1qR monoclonal antibodies on the number of cell-associated *C. albicans* cells. (B) Effects of the anti-gC1qR antibody 74.5.2 and the EGFR kinase inhibitor, gefitinib on the number of cell-associated *C. albicans* cells. (C and D) Number of *C. albicans* cells that are cell-associated with NIH/3T3 cells expressing human gC1qR and/or human EGFR. (C) Effects of EGFR and gC1qR expression on cell-association. (D) Effects of inhibiting surface-expressed gC1qR with the anti-gC1qR antibody 74.5.2. Results are the mean ± SD of three independent experiments, each performed in triplicate. The data were analyzed using one-way analysis of variance with Dunnett’s test for multiple comparisons. ns, not significant; \**P* < 0.05.

**Fig. S3.**
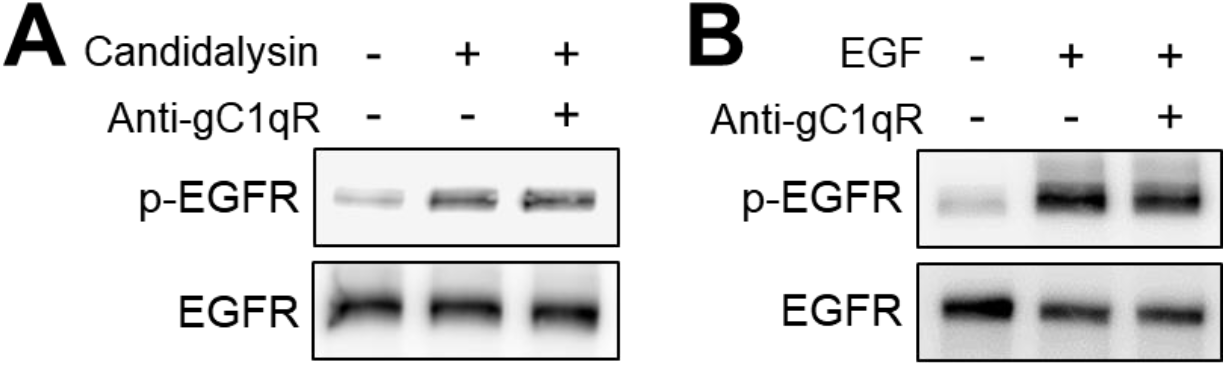
Inhibition of gC1qR does not block EGFR phosphorylation in response to 40 μM candidalysin or 1 ng/ml epidermal growth factor (EGF). Representative immunoblots from three independent experiments.

**Fig. S4.**
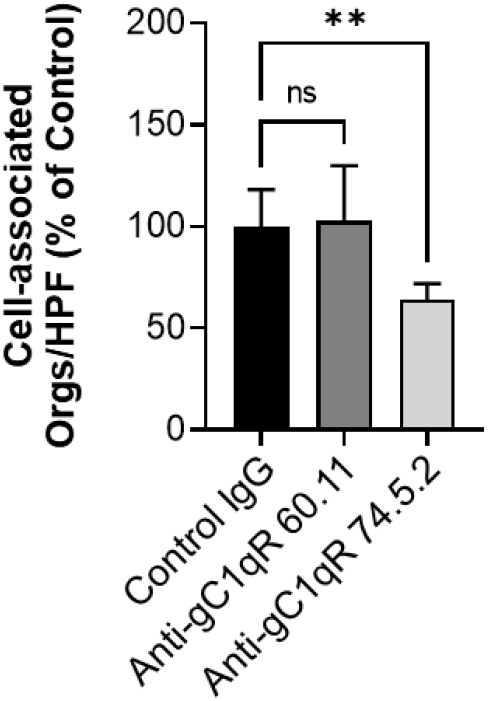
Effects of the indicated anti-gC1qR antibodies on the number of C. albicans cells that were associated with primary mouse oral epithelial cells. Results are the mean ± SD of three independent experiments, each performed in triplicate. The data were analyzed using one-way analysis of variance with Dunnett’s test for multiple comparisons. ns, not significant; \*\**P* < 0.01.

## REFERENCES

1. Hauman CH, Thompson IO, Theunissen F, Wolfaardt P. 1993. Oral carriage of *Candida*in healthy and HIV-seropositive persons. Oral Surg Oral Med Oral Pathol 76:570–2.

2. Verma A, Gaffen SL, Swidergall M. 2017. Innate immunity to mucosal *Candida* infections. J Fungi 3:60.

3. Berberi A, Noujeim Z, Aoun G. 2015. Epidemiology of oropharyngeal candidiasis in human immunodeficiency virus/acquired immune deficiency syndrome patients and CD4+ counts. J Int Oral Health 7:20–3.

4. Gligorov J, Bastit L, Gervais H, Henni M, Kahila W, Lepille D, Luporsi E, Sasso G, Varette C, Azria D. 2011. Prevalence and treatment management of oropharyngeal candidiasis in cancer patients: results of the French CANDIDOSCOPE study. Int J Radiat Oncol Biol Phys 80:532–9.

5. Sanitá PV, Pavarina AC, Giampaolo ET, Silva MM, Mima EG, Ribeiro DG, Vergani CE. 2011. *Candida* spp. prevalence in well controlled type 2 diabetic patients with denture stomatitis. Oral Surg Oral Med Oral Pathol Oral Radiol Endod 111:726–33.

6. Schwartz IS, Patterson TF. 2018. The emerging threat of antifungal resistance in transplant infectious diseases. Curr Infect Dis Rep 20:2.

7. Patel PK, Erlandsen JE, Kirkpatrick WR, Berg DK, Westbrook SD, Louden C, Cornell JE, Thompson GR, Vallor AC, Wickes BL, Wiederhold NP, Redding SW, Patterson TF. 2012. The changing epidemiology of oropharyngeal candidiasis in patients with HIV/AIDS in the era of antiretroviral therapy. AIDS Res Treat 2012:262471.

8. Dotta L, Scomodon O, Padoan R, Timpano S, Plebani A, Soresina A, Lougaris V, Concolino D, Nicoletti A, Giardino G, Licari A, Marseglia G, Pignata C, Tamassia N, Facchetti F, Vairo D, Badolato R. 2016. Clinical heterogeneity of dominant chronic mucocutaneous candidiasis disease: presenting as treatment-resistant candidiasis and chronic lung disease. Clin Immunol 164:1–9.

9. Aggor FEY, Break TJ, Trevejo-Nuñez G, Whibley N, Coleman BM, Bailey RD, Kaplan DH, Naglik JR, Shan W, Shetty AC, McCracken C, Durum SK, Biswas PS, Bruno VM, Kolls JK, Lionakis MS, Gaffen SL. 2020. Oral epithelial IL-22/STAT3 signaling licenses IL-17-mediated immunity to oral mucosal candidiasis. Sci Immunol 5:eaba0570.

10. Conti Heather R, Bruno Vincent M, Childs Erin E, Daugherty S, Hunter Joseph P, Mengesha Bemnet G, Saevig Danielle L, Hendricks Matthew R, Coleman Bianca M, Brane L, Solis N, Cruz JA, Verma Akash H, Garg Abhishek V, Hise Amy G, Richardson Jonathan P, Naglik Julian R, Filler Scott G, Kolls Jay K, Sinha S, Gaffen Sarah L. 2016. IL-17 Receptor signaling in oral epithelial cells is critical for protection against oropharyngeal candidiasis. Cell Host Microbe 20:606–617.

11. Swidergall M, Solis NV, Lionakis MS, Filler SG. 2018. EphA2 is an epithelial cell pattern recognition receptor for fungal b-glucans. Nat Microbiol 3:53–61.

12. Liu Y, Shetty AC, Schwartz JA, Bradford LL, Xu W, Phan QT, Kumari P, Mahurkar A, Mitchell AP, Ravel J, Fraser CM, Filler SG, Bruno VM. 2015. New signaling pathways govern the host response to *C. albicans* infection in various niches. Genome Res 25:679–89.

13. Zhu W, Phan QT, Boontheung P, Solis NV, Loo JA, Filler SG. 2012. EGFR and HER2 receptor kinase signaling mediate epithelial cell invasion by *Candida albicans* during oropharyngeal infection. Proc Natl Acad Sci U S A 109:14194–14199.

14. Solis NV, Swidergall M, Bruno VM, Gaffen SL, Filler SG. 2017. The aryl hydrocarbon receptor governs epithelial cell invasion during oropharyngeal candidiasis. mBio 8: pii: e00025-17.

15. Ho J, Yang X, Nikou SA, Kichik N, Donkin A, Ponde NO, Richardson JP, Gratacap RL, Archambault LS, Zwirner CP, Murciano C, Henley-Smith R, Thavaraj S, Tynan CJ, Gaffen SL, Hube B, Wheeler RT, Moyes DL, Naglik JR. 2019. Candidalysin activates innate epithelial immune responses via epidermal growth factor receptor. Nat Commun 10:2297.

16. Swidergall M, Solis NV, Millet N, Huang MY, Lin J, Phan QT, Lazarus MD, Wang Z, Yeaman MR, Mitchell AP, Filler SG. 2021. Activation of EphA2-EGFR signaling in oral epithelial cells by *Candida albicans* virulence factors. PLoS Pathog 17:e1009221.

17. Dickson MA, Hahn WC, Ino Y, Ronfard V, Wu JY, Weinberg RA, Louis DN, Li FP, Rheinwald JG. 2000. Human keratinocytes that express hTERT and also bypass a p16(INK4a)-enforced mechanism that limits life span become immortal yet retain normal growth and differentiation characteristics. Mol Cell Biol 20:1436–47.

18. Phan QT, Myers CL, Fu Y, Sheppard DC, Yeaman MR, Welch WH, Ibrahim AS, Edwards JE, Filler SG. 2007. Als3 is a *Candida albicans* invasin that binds to cadherins and induces endocytosis by host cells. PLoS Biol 5:e64.

19. El-Sayed A, Harashima H. 2013. Endocytosis of gene delivery vectors: from clathrin-dependent to lipid raft-mediated endocytosis. Mol Ther 21:1118–30.

20. Otto GP, Nichols BJ. 2011. The roles of flotillin microdomains--endocytosis and beyond. J Cell Sci 124:3933–40.

21. Moreno-Ruiz E, Galan-Diez M, Zhu W, Fernandez-Ruiz E, d’Enfert C, Filler SG, Cossart P, Veiga E. 2009. *Candida albicans* internalization by host cells is mediated by a clathrin-dependent mechanism. Cell Microbiol 11:1179–1189.

22. Park H, Myers CL, Sheppard DC, Phan QT, Sanchez AA, Edwards JE, Jr., Filler SG. 2005. Role of the fungal Ras-protein kinase A pathway in governing epithelial cell interactions during oropharyngeal candidiasis. Cell Microbiol 7:499–510.

23. Moreau V, Galan JM, Devilliers G, Haguenauer-Tsapis R, Winsor B. 1997. The yeast actin-related protein Arp2p is required for the internalization step of endocytosis. Mol Biol Cell 8:1361–75.

24. Engqvist-Goldstein AE, Zhang CX, Carreno S, Barroso C, Heuser JE, Drubin DG. 2004. RNAi-mediated Hip1R silencing results in stable association between the endocytic machinery and the actin assembly machinery. Mol Biol Cell 15:1666–79.

25. Phillips DR, Jennings LK, Edwards HH. 1980. Identification of membrane proteins mediating the interaction of human platelets. J Cell Biol 86:77–86.

26. Schaks M, Giannone G, Rottner K. 2019. Actin dynamics in cell migration. Essays Biochem 63:483–495.

27. Itoh RE, Kiyokawa E, Aoki K, Nishioka T, Akiyama T, Matsuda M. 2008. Phosphorylation and activation of the Rac1 and Cdc42 GEF Asef in A431 cells stimulated by EGF. J Cell Sci 121:2635–42.

28. Braun L, Ghebrehiwet B, Cossart P. 2000. gC1q-R/p32, a C1q-binding protein, is a receptor for the InlB invasion protein of *Listeria monocytogenes*. Embo J 19:1458–66.

29. Mengaud J, Ohayon H, Gounon P, Mege RM, Cossart P. 1996. E-cadherin is the receptor for internalin, a surface protein required for entry of L. monocytogenes into epithelial cells. Cell 84:923–32.

30. Veiga E, Cossart P. 2005. *Listeria* hijacks the clathrin-dependent endocytic machinery to invade mammalian cells. Nat Cell Biol 7:894–900.

31. Zatloukal B, Kufferath I, Thueringer A, Landegren U, Zatloukal K, Haybaeck J. 2014. Sensitivity and specificity of in situ proximity ligation for protein interaction analysis in a model of steatohepatitis with Mallory-Denk bodies. PLoS One 9:e96690.

32. Li Z, Lim SK, Liang X, Lim YP. 2018. The transcriptional coactivator WBP2 primes triplenegative breast cancer cells for responses to Wnt signaling via the JNK/Jun kinase pathway. J Biol Chem 293:20014–20028.

33. Moyes DL, Shen C, Murciano C, Runglall M, Richardson JP, Arno M, Aldecoa-Otalora E, Naglik JR. 2014. Protection against epithelial damage during *Candida albicans* infection is mediated by PI3K/Akt and mammalian target of rapamycin signaling. J Infect Dis 209:1816–26.

34. Moyes DL, Runglall M, Murciano C, Shen C, Nayar D, Thavaraj S, Kohli A, Islam A, Mora-Montes H, Challacombe SJ, Naglik JR. 2010. A biphasic innate immune MAPK response discriminates between the yeast and hyphal forms of *Candida albicans* in epithelial cells. Cell Host Microbe 8:225–35.

35. Zhang Y, Lee C, Geng S, Li L. 2019. Enhanced tumor immune surveillance through neutrophil reprogramming due to Tollip deficiency. JCI Insight 4.

36. Shah JA, Vary JC, Chau TT, Bang ND, Yen NT, Farrar JJ, Dunstan SJ, Hawn TR. 2012. Human TOLLIP regulates TLR2 and TLR4 signaling and its polymorphisms are associated with susceptibility to tuberculosis. J Immunol 189:1737–46.

37. Didierlaurent A, Brissoni B, Velin D, Aebi N, Tardivel A, Käslin E, Sirard JC, Angelov G, Tschopp J, Burns K. 2006. Tollip regulates proinflammatory responses to interleukin-1 and lipopolysaccharide. Mol Cell Biol 26:735–42.

38. Spence JS, He R, Hoffmann HH, Das T, Thinon E, Rice CM, Peng T, Chandran K, Hang HC. 2019. IFITM3 directly engages and shuttles incoming virus particles to lysosomes. Nat Chem Biol 15:259–268.

39. Jiang J, Zhang Y, Krainer AR, Xu RM. 1999. Crystal structure of human p32, a doughnut-shaped acidic mitochondrial matrix protein. Proc Natl Acad Sci U S A 96:3572–7.

40. Joseph K, Shibayama Y, Nakazawa Y, Peerschke EI, Ghebrehiwet B, Kaplan AP. 1999. Interaction of factor XII and high molecular weight kininogen with cytokeratin 1 and gC1qR of vascular endothelial cells and with aggregated Abeta protein of Alzheimer’s disease. Immunopharmacology 43:203–10.

41. Lim BL, Reid KB, Ghebrehiwet B, Peerschke EI, Leigh LA, Preissner KT. 1996. The binding protein for globular heads of complement C1q, gC1qR. Functional expression and characterization as a novel vitronectin binding factor. J Biol Chem 271:26739–44.

42. Eggleton P, Ghebrehiwet B, Sastry KN, Coburn JP, Zaner KS, Reid KB, Tauber AI. 1995. Identification of a gC1q-binding protein (gC1q-R) on the surface of human neutrophils. Subcellular localization and binding properties in comparison with the cC1q-R. J Clin Invest 95:1569–78.

43. Ghebrehiwet B, Lim BL, Peerschke EI, Willis AC, Reid KB. 1994. Isolation, cDNA cloning, and overexpression of a 33-kD cell surface glycoprotein that binds to the globular “heads” of C1q. J Exp Med 179:1809–21.

44. Peerschke EI, Bayer aS, Ghebrehiwet B, Xiong YQ. 2006. gC1qR/p33 blockade reduces *Staphylococcus aureus* colonization of target tissues in an animal model of infective endocarditis. Infect Immun 74:4418–23.

45. Biswas AK, Hafiz A, Banerjee B, Kim KS, Datta K, Chitnis CE. 2007. *Plasmodium falciparum* uses gC1qR/HABP1/p32 as a receptor to bind to vascular endothelium and for platelet-mediated clumping. PLoS Pathog 3:1271–80.

46. Kim BC, Hwang HJ, An HT, Lee H, Park JS, Hong J, Ko J, Kim C, Lee JS, Ko YG. 2016. Antibody neutralization of cell-surface gC1qR/HABP1/SF2-p32 prevents lamellipodia formation and tumorigenesis. Oncotarget 7:49972–49985.

47. Ghebrehiwet B, Lu PD, Zhang W, Lim BL, Eggleton P, Leigh LE, Reid KB, Peerschke EI. 1996. Identification of functional domains on gC1Q-R, a cell surface protein that binds to the globular “heads” of C1Q, using monoclonal antibodies and synthetic peptides. Hybridoma 15:333–42.

48. Gaborit N, Larbouret C, Vallaghe J, Peyrusson F, Bascoul-Mollevi C, Crapez E, Azria D, Chardes T, Poul MA, Mathis G, Bazin H, Pelegrin A. 2011. Time-resolved fluorescence resonance energy transfer (TR-FRET) to analyze the disruption of EGFR/HER2 dimers: a new method to evaluate the efficiency of targeted therapy using monoclonal antibodies. J Biol Chem 286:11337–45.

49. Moyes DL, Wilson D, Richardson JP, Mogavero S, Tang SX, Wernecke J, Hofs S, Gratacap RL, Robbins J, Runglall M, Murciano C, Blagojevic M, Thavaraj S, Forster TM, Hebecker B, Kasper L, Vizcay G, Iancu SI, Kichik N, Hader A, Kurzai O, Luo T, Kruger T, Kniemeyer O, Cota E, Bader O, Wheeler RT, Gutsmann T, Hube B, Naglik JR. 2016. Candidalysin is a fungal peptide toxin critical for mucosal infection. Nature 532:64–8.

50. Riise GC, Andersson BA, Kjellstrom C, Martensson G, Nilsson FN, Ryd W, Schersten H. 1999. Persistent high BAL fluid granulocyte activation marker levels as early indicators of bronchiolitis obliterans after lung transplant. Eur Respir J 14:1123–30.

51. Schultz J, Kaminker K. 1962. Myeloperoxidase of the leucocyte of normal human blood. I. Content and localization. Arch Biochem Biophys 96:465–467.

52. Gouin E, Gantelet H, Egile C, Lasa I, Ohayon H, Villiers V, Gounon P, Sansonetti PJ, Cossart P. 1999. A comparative study of the actin-based motilities of the pathogenic bacteria *Listeria monocytogenes, Shigella flexneri* and *Rickettsia conorii*. J Cell Sci 112 (Pt 11):1697–708.

53. Skoble J, Auerbuch V, Goley ED, Welch MD, Portnoy DA. 2001. Pivotal role of VASP in Arp2/3 complex-mediated actin nucleation, actin branch-formation, and Listeria monocytogenes motility. J Cell Biol 155:89–100.

54. Rodnick-Smith M, Luan Q, Liu SL, Nolen BJ. 2016. Role and structural mechanism of WASP-triggered conformational changes in branched actin filament nucleation by Arp2/3 complex. Proc Natl Acad Sci U S A 113:E3834–43.

55. Song X, Yao Z, Yang J, Zhang Z, Deng Y, Li M, Ma C, Yang L, Gao X, Li W, Liu J, Wei L. 2016. HCV core protein binds to gC1qR to induce A20 expression and inhibit cytokine production through MAPKs and NF-κB signaling pathways. Oncotarget 7:33796–808.

56. Sethi S, Herrmann M, Roller J, von Müller L, Peerschke EI, Ghebrehiwet B, Bajric I, Menger MD, Laschke MW. 2011. Blockade of gC1qR/p33, a receptor for C1q, inhibits adherence of *Staphylococcus aureus* to the microvascular endothelium. Microvasc Res 82:66–72.

57. Peerschke EI, Ghebrehiwet B. 2007. The contribution of gC1qR/p33 in infection and inflammation. Immunobiology 212:333–42.

58. Shen Y, Naujokas M, Park M, Ireton K. 2000. InIB-dependent internalization of *Listeria* is mediated by the Met receptor tyrosine kinase. Cell 103:501–10.

59. Khelef N, Lecuit M, Bierne H, Cossart P. 2006. Species specificity of the *Listeria monocytogenes* InlB protein. Cell Microbiol 8:457–70.

60. Liu Y, Mittal R, Solis NV, Prasadarao NV, Filler SG. 2011. Mechanisms of *Candida albicans* trafficking to the brain. PLoS Pathog 7:e1002305.

61. Phan QT, Eng DK, Mostowy S, Park H, Cossart P, Filler SG. 2013. Role of endothelial cell septin 7 in the endocytosis of *Candida albicans*. mBio 4:00542–13.

62. Solis NV, Filler SG. 2012. Mouse model of oropharyngeal candidiasis. Nature protocols 7:637–42.

